# Vesicular Rps6 released by astrocytes regulate local translation and enhance synaptic markers in neurons

**DOI:** 10.1101/2025.05.28.656644

**Authors:** María Gamarra, Aida de la Cruz-Gambra, Maite Blanco-Urrejola, Esperanza González, Mikel Azkargorta, Felix Elortza, Juan Manuel Falcón-Pérez, Jimena Baleriola

**Affiliations:** Achucarro Basque Center for Neuroscience, 48940 Leioa, Spain; Departamento de Neurociencias, Universidad del País Vasco (UPV/EHU), 48940 Leioa, Spain; Center for Cooperative Research in Biosciences (CIC bioGUNE), Basque Research and Technology Alliance (BRTA), 48160 Derio, Bizkaia, Spain; Centro de Investigación Biomédica en Red: Enfermedades Hepáticas y Digestivas (CIBERehd), Madrid, Spain; IKERBASQUE Basque Foundation for Science, 48009 Bilbao, Spain

**Keywords:** astroglia-derives EVs, amyloid pathology, intra-axonal protein synthesis, ribosomal proteins, synapses

## Abstract

In neurons, like in any other cell, their function often relies on the fine tuning of their protein levels, which is achieved by the balance between protein synthesis and turnover. Defects in protein homeostasis frequently leads to neuronal dysfunction and neurological disorders. Given their extreme morphological complexity and high compartmentalization, neurons highly depend on the asymmetrical distribution of their proteome. The common belief is that proteins that sustain axonal, dendritic and synaptic functions are synthesized in the soma and then transported to distal neuronal compartments. However, there is a complementary mechanism by which the mRNAs, and not the proteins, are transported to distal subneuronal domains, and once they reach their destination they are locally translated. Although once considered heretical, local translation (or local protein synthesis) is now widely accepted by the scientific community. Nonetheless there is one question that remains largely unexplored in the field and that is whether local translation in dendrites, axons and synapses is fully regulated by the neuron itself or if non-neuronal cells (e.g. glia) can modulate this mechanism in a non-cell-autonomous manner. Here we show that astroglia regulates local protein synthesis and enhances synaptic markers by releasing extracellular vesicles (EVs) containing ribosomal protein Rps6. To our knowledge this is the first report that directly demonstrates glial control of local translation in neurons through EV-mediated glia-to-neuron communication.

## Introduction

Extracellular vesicles (EVs) are membranous nanoparticles secreted by virtually all cell types. EVs can be detected in human fluids by non-invasive methods, and their content closely mirrors the progression of several diseases ^1^. Thus, over the last years EVs have emerged as valuable tools for biomarker discovery and diagnosis. More importantly, due to their low immunogenicity and their high lipid content EVs can be used as potential vehicles to target different organs, including the brain, with therapeutic molecules ^2^. EVs have been recently recognized as relevant players in intercellular communication, as they can transfer proteins, nucleic acids and lipids to acceptor cells where these bioactive molecules elicit their function ^3^. This work was driven by the need to expand the knowledge on the role and applicability EVs to neuronal function and dysfunction. Some authors reported the relevance of EVs in neuronal survival ^4,5^, synaptic plasticity ^6–8^, or neuroinflammation ^9–11^. Given the complexity of neuronal architecture and the sensitivity of the brain to disruptions in intercellular signaling, understanding how EVs function in the central nervous system (CNS) may reveal novel mechanisms involved in both health and disease.

Neurons are the most polarized cells in the brain and their function heavily relies on the accurate asymmetrical distribution of proteins, some of them being enriched in the soma and others in dendrites, axons or synaptic compartments. Local proteomes in neurons can be shaped by proteins synthesized in the soma and then transported to subneuronal domains or by localized mRNAs that are then translated at target sites. This latter mechanism, known as local translation or local protein synthesis, enables subcellular compartments to react to environmental changes in an acute manner, as proteins are produced at limiting amounts in sites where they are needed and only when they are needed. Local translation has been best studied in physiological conditions, where it is involved in axon pathfinding, synapse formation, synaptic plasticity and axonal maintenance to mention but a few ^12–15^. However, recent evidence suggests this mechanism is dysregulated in neurodegenerative and neurological disorders and might contribute to disease progression. For instance, exogenous application of Aβ peptides, major drivers of amyloid pathology in Alzheimer’s disease (AD), induces local protein synthesis in axons, leading to transcriptional changes involved in neurodegeneration ^16,17^. Nevertheless, despite the growing knowledge of local translation in both neuronal physiology or pathology, the contribution of non-neuronal cells to this mechanism has remained elusive. Whereas the neuronal soma is considered the primary origin of axonal RNAs, increasing hypotheses suggest that surrounding glia might be an additional source of mRNAs and ribosomes localized to axons. Indeed, several evidence indicates the existence of active RNA and ribosomal protein transfer from Schwann cells to regenerating nerves ^18,19^. But perhaps the most striking observation regarding the potential role of glia in local axonal translation was recently published by Müller *et al*.: using two glial RiboTracker Cre mouse lines, the authors found that ribosomes appearing in axons following sciatic nerve injury were predominantly glial ^20^. A few mechanisms have been proposed for RNA and ribosome transfer from glia to axons, but these and other bioactive molecules are likely delivered to axons within EVs. Notably, Shwann cell-derived EVs improve regeneration of injured peripheral nerves ^21^. In this context, it has been proposed that glial-EVs induce regeneration by changing the axonal translatome in the peripheral nervous system (PNS). However, there is no direct evidence on the regulation of local neuronal translation by glial EVs in the central nervous system (CNS).

Here we investigated whether neuron-astrocyte communication through secreted factors regulated local translation in neurons. We identified astroglia EVs released upon Aβ treatment as positive modulators of intra-axonal protein synthesis. Interestingly, these EVs also enhanced the levels of synaptic markers. Finally, proteomic analyses of astrocytic-derived EV cargoes revealed that ribosomal protein Rps6 secreted by glia was responsible for EV-mediated local protein synthesis, and its genetic downregulation led to a decrease in synaptic proteins. Overall, our results suggest that astroglia EVs released in response to Aβ contain ribosomal proteins that regulate local translation in neurons through non-cell-autonomous mechanisms. In turn, locally synthesized proteins enhance synaptic markers potentially contributing to synaptic function.

## Materials and Methods

### 1. Animals

All animal procedures were conducted in accordance with the European Directive 2010/63/EU and were approved by the University of the Basque Country (UPV/EHU) Ethics Committee. Sprague-Dawley rats were bred in local facilities. Embryonic brains were obtained from CO_2_-euthanized pregnant rats for neuronal cultures, while postnatal brains were used for astroglial cultures.

### 2. Primary neuronal cultures

#### 2.1 Cultures in multi-well plates

Hippocampal neurons were harvested from Sprague-Dawley embryonic day 18 (E18) rat embryos as previously described ^22^. Briefly, hippocampal tissue was dissected in ice-cold 1X Hank’s Balanced Salt Solution (HBSS; Gibco, Thermo Fisher Scientific, Waltham MA, USA) and dissociated in TrypLE Express (Gibco) for 15 min at 37 °C in a 5% CO_2_ humidified incubator, followed by mechanical homogenization. Cells were centrifuged at 200 x g for 5 min at room temperature (RT) and resuspended in plating medium containing 10 % fetal bovine serum (FBS; Sigma-Aldrich Aldrich, Merck, Darmstadt, Germany), 2 mM L-glutamine and a mixture of antibiotics (50 U/μL penicillin and 50 μg/μL streptomycin) in Neurobasal medium (all from Gibco). Hippocampal neurons were cultured on multiwell plates pre-coated with poly-D-lysine (PDL; Sigma-Aldrich, #P1149) at different densities depending on the experimental approach: 15.000 cells/cm^2^ for immunofluorescence assays, and 25.000 cells/cm^2^ for protein extraction and EVs isolation. Cultures were maintained at 37 °C in a 5% CO_2_ humidified incubator. At 1 day in vitro (1 DIV), the plating medium was completely replaced for growth medium containing 1X B27 (Gibco), 2 mM L-glutamine and the antibiotic mixture in Neurobasal medium. To restrict glial proliferation, the growth medium was supplemented with the antimitotic agents 5-fluorodeoxyuridine (FdU) and uridine (Sigma-Aldrich), both at 20 μM. Half of the medium was refreshed every 3 days with growth medium containing 20 μM FdU and 20 μM uridine. Treatments were performed at 9-10 DIV.

#### 2.2 Cultures in modified Boyden chambers

To allow neurite outgrowth while preventing soma migration, neurons were cultured in modified Boyden chambers, which consist of a 1-μm pore polyethylene terephthalate membrane (Falcon, Corning, Corning NY, USA). This system also enables neuronal co-culture with astrocytes without direct contact, facilitating the study of cell-cell communication through secreted factors.

Cortical and hippocampal neurons were plated in PDL-coated Boyden chambers at a density of 500.000 cells/cm^2^ in plating medium. Cultures were maintained at 37 °C in a 5% CO_2_ humidified incubator. At 1 DIV, the medium was fully replaced with growth medium supplemented with 20 μM FdU and 20 μM uridine, and half of the medium was refreshed every 3 days. Treatments were performed at 9-10 DIV.

#### 2.3 Cultures in microfluidic chambers

Microfluidic chambers are two- or three-compartment polydimethylsiloxane (PDMS; Dow, Midland MI, USA) devices that enable spatial isolation of different neuronal regions. PDMS was prepared in a 9:1 ratio (polymer-to-curing agent), cast into molds, and cured overnight at 60 °C. Devices were then punched and sterilized with 70 % ethanol and ultraviolet light exposure. Finally, chambers were placed on 30 mm coverslips for immunofluorescence assays.

Hippocampal neurons were seeded in the proximal compartment of PDL-coated microfluidic devices at a density of 100.000 – 150.000 cells per chamber in plating medium. At 1 DIV, the medium was fully replaced with growth medium supplemented with 20 μM FdU and 20 μM uridine. When required, distal compartment medium was enriched with 2X B27 to promote axonal growth. Treatments were performed once optimal axonal confluence was reached.

### 3. Primary astroglia cultures

Astrocyte cultures were obtained from mixed glial cultures following the protocol described by McCarthy and de Vellis (1980) ^23^ with minor modifications. Briefly, cortico-hippocampal tissue was isolated from Sprague-Dawley postnatal day 0–2 (P0–P2) rat pups and dissociated in 0.25% trypsin (Sigma-Aldrich) and 0.004% DNase (Sigma-Aldrich) for 15 min at 37 °C in a 5% CO₂ humidified incubator. The enzymatic reaction was stopped by adding plating medium containing 10 % FBS (HyClone; Cytiva Life Science, Marlborough MA, USA), 50 U/mL penicillin-streptomycin and 250 ng/mL amphotericin G (Gibco) in Iscove’s Modified Dulbecco’s Medium (IMDM; Gibco). Cells were centrifuged at 300 x g for 6 min at RT, resuspended and mechanically dissociated using 21G and 23G needles, respectively. The cell suspension was centrifuged again under the same conditions and resuspended in plating medium. Mixed glial cells were then seeded onto 75 cm^2^ PDL (Sigma-Aldrich, #P6407)-coated flasks (BioLite, Thermo Fisher Scientific) and maintained at 37 °C in a 5% CO_2_ humidified incubator. At 1 DIV, the plating medium was replaced with growth medium consisting of Dulbecco’s Modified Eagle Medium (DMEM; Gibco) supplemented with 10 % FBS (Sigma-Aldrich), 2 mM L-glutamine and 50 U/mL penicillin-streptomycin. The medium was fully exchanged every 3 days.

To isolate astrocytes from mixed glial cultures, flasks at 11 DIV were shaken at 180 rpm for 4 h at 37 °C, allowing the removal of microglia while astrocytes remained adhered to the flask surface. Astrocytes were then detached by incubation with TrypLE Express for 15 min at 37 °C in a 5% CO_2_ humidified incubator. The enzymatic reaction was stopped by adding medium containing 10% FBS and cells were centrifuged at 300 x g for 5 min at RT. The astrocyte-containing pellet was resuspended in neuronal growth medium.

### 4. Neuron-astrocyte co-cultures

At 11 DIV, astrocytes were collected as described in the previous section. For modified Boyden chambers, astrocytes were seeded at the bottom of 7 DIV neuronal cultures in a 1:10 astrocyte-to-neuron ratio. For validation of proteomic analyses by immunocytochemistry, to maintain indirect communication between neurons and astrocytes, neurons were cultured on coverslips, while astrocytes were plated on the lower membrane side of modified Boyden chambers. For neuron-astrocytes co-cultures in microfluidic chambers, astroglia were cultured in the axonal compartment (once axons had reached confluency) in a 1:10 astrocyte-to-neuron ratio. In this case, however, both neurons and astrocytes were isolated from E17.5 mouse embryos.

### 5. Aβ oligomerization and treatment

Soluble oligomeric amyloid-β_1-42_ (Aβ_1-42_) was prepared as previously described ^24^. Synthetic human Aβ_1-42_ (Bachem, Bubendorf, Switzerland) was dissolved in hexafluoroisopropanol (HFIP; Sigma-Aldrich) at 1 mM, aliquoted, and dried. Peptides were then resuspended in 5 mM dimethyl sulfoxide (DMSO; Sigma-Aldrich) and diluted in Ham’s F-12 medium (pH 7.4; PromoCell, Labclinics, Barcelona, Spain) to a final concentration of 100 μM. To induce oligomerization, Aβ peptides were incubated overnight at 4 °C.

At 9 – 10 DIV neurons were treated with 3 μM oligomeric Aβ in neuronal growth medium. Treatment of astrocytes was performed in 2-3 DIV cultures after astroglia were isolated from mixed glial cultures (11 DIV in flask + 2-3 DIV in plates). In co-cultures in modified Boyden, the treatment was applied to the lower compartment. All treatments were performed for 24 hours and DMSO was used as vehicle control.

### 6. Detection of newly synthesized proteins by puromycin labelling

#### 6.1 Puromycilation assay followed by immunocytochemistry

Puromycin is a structural analog of the aminoacyl-tRNA that carries tyrosine. During active translation elongation, puromycin/it incorporates into the nascent polypeptide in a ribosome catalyzed reaction ^25^. Nascent peptides can then be identified with anti-puromycin antibodies. For this assays, neuronal cultures and neuron-astrocytes co-cultures, previously treated with vehicle or Aβ, were incubated with 2 μM puromycin (Sigma-Aldrich) for 30 min at 37 °C. In the case of neurons grown in microfluidic chambers, puromycin was specifically applied to the axonal compartment. After incubation, cells were washed with 1X phosphate-buffered saline (PBS; pH 7.4) containing 3 μg/mL digitonin (Sigma-Aldrich) and fixed in 4 % paraformaldehyde (PFA; Fisher Chemical, Pittsburgh PA, USA) with 4 % sucrose. Puromycin incorporation was analyzed by immunocytochemistry.

#### 6.2 Click chemistry

Alternatively, newly synthesized proteins were detected with O-propargyl-puromycin (OPP). OPP is a modified version of puromycin that contains an alkyne group, which allows for a covalent reaction with an azide-conjugated biotin. This biotinylation step enhances detection sensitivity while minimizing background signal ^26^. To study *de novo* synthesis of neuritic proteins, 24 h after vehicle or Aβ treatment, neuronal and neuron-astrocyte co-cultures in modified Boyden chambers were incubated with 25 µM OPP (Jena Bioscience, Jena, Germany) in the lower compartment for 1 h at 37 °C. After incubation, the somatic region was scraped out with a cotton swab and the neuritic fraction was washed with PBS. Proteins were extracted using a lysis buffer containing 20 mM HEPES (Fisher Bioreagent, Pittsburgh PA, USA), 100 mM sodium ascorbate (Jena Bioscience) and 5 mM magnesium acetate (pH 7.5; Alfa Aesar, Thermo Fisher Scientific). The click chemistry reaction was performed by conjugating the OPP alkyne group to azide-PEG3-biotine (Jena Bioscience) in a Cu(I)-catalyzed reaction. Biotinylated proteins were then precipitated with acetone for 2 h at −20 °C, centrifuged at 15,000 x g for 5 min at RT, washed with methanol and sonicated for 5 s. Proteins were centrifuged at 15,000 x g for 2 min at RT, and again washed with methanol and sonicated for 5 s. This process was repeated twice. The final protein pellet was resuspended in a loading buffer containing LDS (pH 8.5), SERVA Blue G250 and Red Phenol (Invitrogen) and 1,4-dithiothreitol (DTT; Thermo Fisher Scientific). Biotinylated proteins were detected by western blotting using streptavidin.

### 7. Extracellular vesicle (EV) isolation and labelling

EVs were isolated from conditioned media of neuronal or astroglia cultures. After the corresponding treatment, culture media were collected, and cell debris was removed by centrifugation at 200 x g for 5 min at 4 °C. The supernatant was then ultracentrifuged at 250,000 x g for 75 min at 4 °C using an OptimaMAX-XP tabletop ultracentrifuge with a TLA 120.2 rotor (Beckman Coulter, Brea CA, USA). The pellet was then resuspended in 1X PBS to obtain the vesicular fraction. When required, EVs were labelled with the red lipophilic dye PKH26 (Sigma-Aldrich). After ultracentrifugation, the EV pellet was resuspended in 1X PBS and incubated with PKH26 and Diluent C for 5 min at RT. The reaction was stopped by adding 1 % bovine serum albumin (BSA; Thermo Fisher Scientific) in 1X PBS. Labeled EVs were then ultracentrifuged at 250,000 x g for 75 min at 4 °C, as previously described. The final EV pellet was resuspended in 15 μL of 1X PBS. As a negative control, PBS aliquots without EVs were labeled following the aforementioned protocol.

### 8. Treatments with EVs or EV-depleted media

To investigate the role of EVs on neurons, EVs were isolated from neuronal and astroglia cultures, as described above. The volume of EVs for treatments was adjusted to the number of donor cells, previously quantified under an optical microscope. To treat neurons cultured in multi-well plates, the medium was replaced with fresh medium containing EVs and incubated for 45 min at 37 °C in a 5 % CO_2_ humidified incubator. Non-EV supplemented fresh medium was used as control. In microfluidic chambers, EVs were directly applied to the distal axonal compartment. For synaptic marker analyses, EVs were added directly to neuronal cultures and incubated for 24 h at 37 °C. Non-EV supplemented medium was used as a control.

Whenever stated, the effect of EVs was addressed by treating neurons with conditioned media released by astrocytes resulting either from the downregulation of proteins involved in EV biogenesis or from EV depletion by ultracentrifugation. Neuronal EV-depleted medium was also used in selected experiments.

### 9. siRNAs transfection

Astrocytes were transfected with small interfering RNAs (siRNAs) to inhibit target protein expression. Three days before transfection, cells were seeded in Neurobasal medium growth containing 1X B27 (Gibco), 2 mM L-glutamine without antibiotics. Transfection was carried out using 10 nM siRNAs (including a control siRNA; all from Ambion, Thermo Fisher Scientific) and Lipofectamine RNAiMAX (Invitrogen, Thermo Fisher Scientific), following manufacturer’s instructions. Sequences of targeting siRNAs were the following::

- siRNA *Rab27a* (s132277): 5’ – CGCGUACUGCGAAAACCCAtt – 3’
- siRNA *Rab35* (s143711): 5’ – GCAGCUUGCUGUUACGAUUtt – 3’
- siRNA *Rab11a* (s135850): 5’ – GAGAUAUACCGCAUUGUUUtt – 3’
- siRNA *Rps6* (s131129): 5’ – GCAGAAUAUGCUAAACUUUtt – 3’

To address the efficiency of each siRNA on their specific targets, protein and mRNA detection were addressed by western blotting and quantitative RT-PCR 24 hours after transfection. To evaluate whether siRNAs affected EV release, the culture medium was fully replaced 24 h after transfection, and astrocytes were incubated for another 24 h. Conditioned media were then collected for EV isolation and protein detection.

### 10. Immunocitochemistry

After the corresponding treatments, cultures were fixed using 4 % PFA with 4 % sucrose for 20 minutes at RT. Fixed cells were then permeabilized and blocked using 100 mM glycine, 0.25 % Triton X-100 (Thermo Fisher Scientific) and 3 % BSA in 1X PBS. Cells were incubated overnight at 4 °C with primary antibodies (Table I). After 3 washing steps with 1X PBS, cells were incubated for 1 h with the appropriate fluorophore-conjugated secondary antibodies at RT. Samples were washed 3 times with PBS, once with distilled water and mounted using Antifade ProLong Gold with DAPI (Invitrogen).

**Table I:** Primary antibodies used for immunocytochemistry.

| Target | Species | Reference | Dilution |
| --- | --- | --- | --- |
| Puromycin | Mouse monoclonal | MABE343; Merck | 1/500 |
| Tau | Rabbit monoclonal | ab32057, Abcam | 1/500 |
| Map-2 | Chicken polyclonal | AB5543; Merck | 1/1,000 |
| Rpl26 | Rabbit polyclonal | ab59567; Abcam | 1/200 |
| Rps6 | Rabbit polyclonal | ab40820; Abcam | 1/200 |
| Synaptophysin | Chicken polyclonal | 101006; Synaptic Systems | 1/500 |
| Homer-1 | Guinea pig polyclonal | 160005; Synaptic Systems | 1/500 |

### 11. Western blotting

Western blot analyses were used for 1) the detection of nuclear and cytoskeletal proteins in whole cell and neuritic lysates; 2) the detection of biotinylated OPP-labeled proteins extracted from neuritic fractions; 3) the identification of EV markers in cell lysates and vesicular extracts; and 3) the evaluation of Rps6 levels in astrocytes after siRNA transfection.

Whole lysate proteins, neuritic proteins and vesicular proteins were extracted using RIPA buffer supplemented with 1X protease and phosphatase inhibitors (Thermo Fisher Scientific). OPP labelled proteins were isolated with 20 mM HEPES supplemented with 100 mM sodium ascorbate and 5 mM magnesium acetate (pH 7.5). In all case proteins were separated by PAGE-SDS electrophoresis in 4-12 % or 8-16 % polyacrylamide continuous gradient gels in reducing conditions, except for vesicular extracts in which non-reducing conditions were used.

Proteins were transferred onto nitrocellulose membranes, or polyvinylidene difluoride membrane (PVDF; Amersham, Sigma-Aldrich) previously activated with methanol. Membranes were blocked using 5 % BSA in TRIS-buffered saline with 0.1 % Tween (TBS-T; pH 7.6) for 1 h at RT. For EV fractions, 5 % nonfat dried milk in TBS-T was used.

Biotinylated OPP-labeled proteins were detected using HRP-conjugated streptavidin (1:10,000 in 3% BSA in TBS-T) for 2 h at RT. For other proteins, membranes were incubated overnight at 4 °C in agitation with primary antibodies (Table II). After 3 washes with TBS-T, membranes were incubated the appropriate HRP-conjugated secondary antibodies for 1 h at RT. Detection was performed with the SuperSignal™ West Pico PLUS kit (Thermo Fisher Scientific) and imaged with the ChemiDoc MP system (Bio-Rad, Hercules CA, USA). For biotinylated OPP-labeled proteins, membranes were stained with Amido Black for 1 min, washed with 1% acetic acid, dried, and then imaged using ChemiDoc MP. Blots were analyzed with FIJI/ImageJ.

**Table II:**
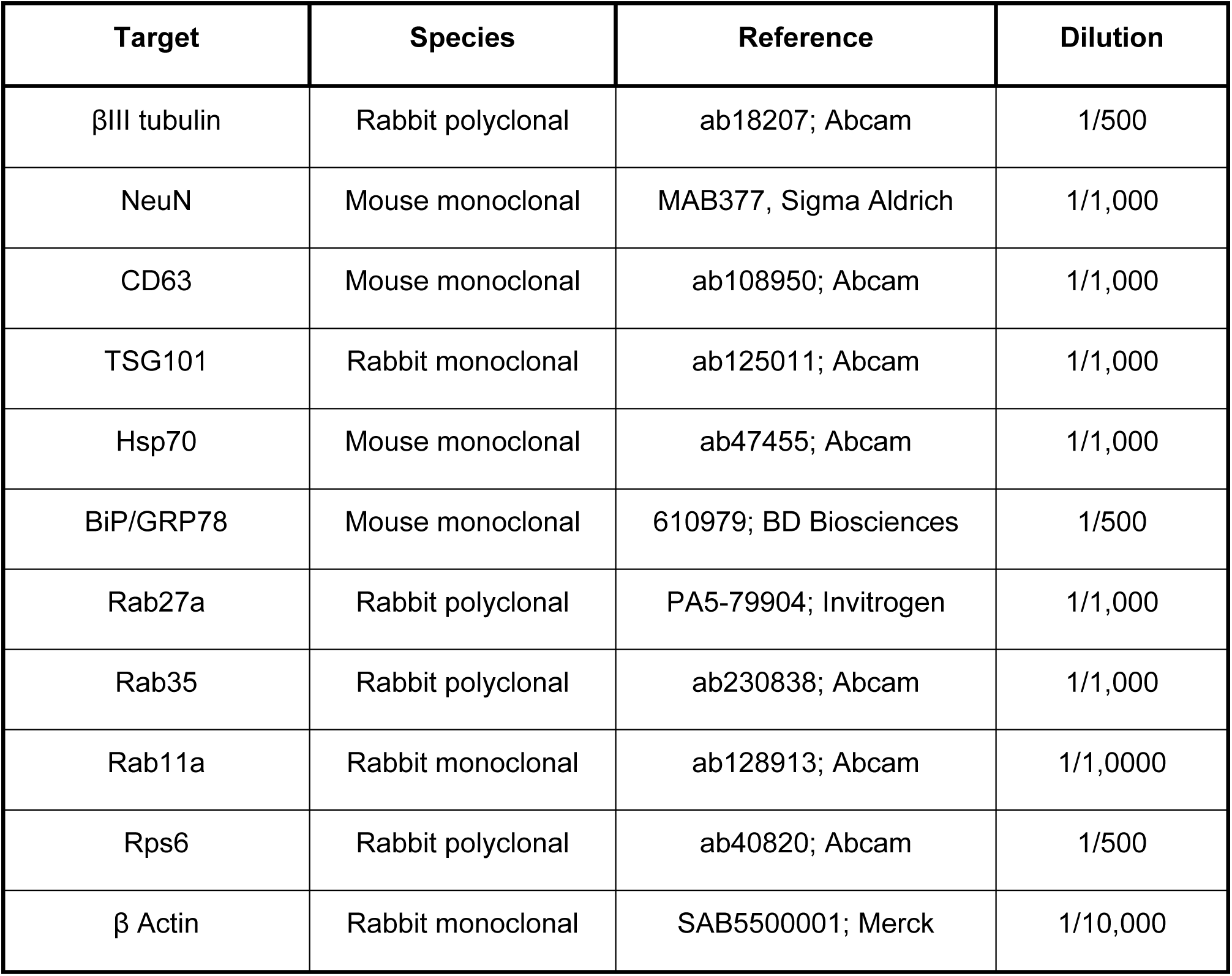
Primary antibodies used for western blotting.

### 12. Real-time RT-PCR

Real-time RT-PCR was used to assess siRNA transfection efficiency. Total RNA was extracted from astrocytes 24 hours post-transfection using Trizol (Thermo Fisher Scientific) following the manufacturer’s instructions. To improve RNA purity, 300–500 ng of RNA was treated with DNase (Invitrogen). RNA concentration was measured with a NanoDrop (Thermo Fisher Scientific). Reverse transcription was performed using the SuperScript IV First-Strand Synthesis System (Invitrogen) with oligo(dT)_20_. cDNA was amplified by real-time PCR using the Power SYBR Green Master Mix (Thermo Fisher Scientific) under the following conditions: 40 cycles of 95 °C for 10 min; 95 °C for 15 s; and 60 °C for 1 min. Primer sequences (forward, f; and reverse, r) are detailed below (Invitrogen; Sigma-Aldrich). Actb and Gapdh were used as housekeeping genes.

*Rab27a* (*Rattus novergicus*): f 5’ – GGGCAGGAGAGGTTTCGTAG – 3’; r: 5’ – TACGCGTGCATCTGTAGC – 3’
*Rab35* (*Rattus novergicus*): f 5’ – ACAACACCTTCTCAGCAGCT – 3’; r 5’ – CTGTCCCAGATCTGCAGCTT – 3’
*Rab11a* (*Rattus novergicus*): f 5’ – ACGTCTGCATACTATCGTGGA – 3’; r 5’ – GAACTGCCCTGAGATGACGT – 3’
*Actb* (*Rattus novergicus*): f 5’ – TACAACCTTCTTGCAGCTCC – 3’; r 5’ – ATACCCACCATCACACCCTG – 3’
*Gapdh* (*Rattus novergicus*): f 5’ – TGCCAGCCTCGTCTCATAGA – 3’; r 5’ – TGACTGTGCCGTTGAACTTG – 3’

### 13. Proteomic analyses by mass spectrometry

For proteome analysis, neurons were cultured in modified Boyden chambers as described previously. After 24 h of DMSO or Aβ treatment, protein content was obtained from neurites or the somatic region by selectively scraping out the undesired fraction and washing the compartment of interest with 1X PBS. Proteins were extracted using a lysis buffer containing 7 M urea (Fisher Chemical), 2 M tiourea (Alfa Aesar), 4 % CHAPS (Acros Organics, Geel, Belgium) and 5 mM DTT in water. For EV proteomic analysis, neuronal- and astroglial-derived EVs were isolated by ultracentrifugation and resuspended in the same buffer. Fresh medium samples were included to account for the potential bovine-derived protein contaminants.

Mass spectrometry was performed in the Proteomics facility from CIC bioGUNE (Derio, Spain). Protein digestion followed the FASP method ^27^, with overnight trypsinization at 37 °C. Peptides were desalted using C18 tips (Millipore, Burlington MA, USA) and analyzed by LC-MS/MS in a timsTOF Pro with PASEF spectrometer (Bruker Daltonics, Billerica MA, USA) connected to an Evosep ONE chromatograph (Evosep, Odense, Denmark). A total of 200 ng per sample were analyzed using the 30 SPD method (∼44 min gradients). Protein identification and quantification were performed using MaxQuant (Tyanova et al., 2016a) with the Uniprot *Rattus norvegicus* proteome as reference. Precursor and fragment mass tolerances were set at 20 ppm and 0.05 Da, respectively. Only proteins with at least two peptides and FDR <1% were considered for further analysis. R ProBatch was used to correct variability, and differential expression analysis was performed using Perseus.

Gene Ontology (GO) analysis was conducted using DAVID software ^28,29^ to determine associated cellular compartments, biological processes, and KEGG pathways (p < 0.05; ≥2 proteins per term). STRING ^30^ was used to analyze protein-protein interaction networks (Medium stringency; FDR < 0.05; MCL= 1.4).

### 14. Cryo-electron microscopy

Neuronal and astroglial-derived EVs were isolated as previously described. Cryo-electron microscopy (Cryo-EM) was performed in the Electron Microscopy and Crystallography facility from CICbioGUNE (Derio, Spain). Samples were adsorbed onto glow-discharged R2/1 300-mesh holey carbon grids (Quantifoil, Jena, Germany), blotted at 95 % humidity, and rapidly vitrified in liquid ethane using a LEICA EM GP2 (Leica, Wetzlar, Germany). Cryo-electron microscopy was performed at liquid nitrogen temperature on a JEM-2200FS/CR transmission electron microscope (JEOL, Tokyo, Japan) equipped with a field emission gun and operated at 200 kV.

### 15. Nanoparticle Tracking Analyisis, NTA

Nanoparticle Tracking Analysis (NTA) was performed using a NanoSight LM10 system (NanoSight, Malvern, UK) with fast video capture and particle-tracking software. Consistent acquisition settings were maintained across all samples in a given experiment. Vesicle size distribution was analyzed based on brownian motion, and the mean and mode of vesicle size were used to estimate particle concentration. In the case of siRNA transfected cells, fresh medium was also subjected to NTA analyses to subtract the noise signal that originated from residual lipofectamine. For all experiments, particle release was normalized to the number of donor cells.

### 16. Image acquisition and processing

#### 16.1. Immunocitochemistry image acquisition

Images were acquired using an Axio-Observer Z1 microscope (Zeiss, Oberkochen, Germany) with Plan-Apochromat 20x/0.8 M27, EC Plan-Neofluar 40x/1.30 Oil DIC M27, and Plan-Apochromat 63x/1.4 Oil DIC M27 objectives (Zeiss). Settings were established in a random field of a control sample, avoiding pixel saturation, and maintained across all samples in each experiment. When possible, images were taken from five random fields per coverslip and two coverslips per experimental condition. Most images were captured with an AxioCam MRm Rev.3 camera (Zeiss), while a Hamamatsu EM-CCD ImagEM camera (Hamamatsu Photonics, Hamamatsu, Japan) was used for the far-red spectrum.

#### 16.2. Fluorescence Intensity analyses in non-binarized Images

Fluorescence intensity was quantified using FIJI/ImageJ, following the method described in Gamarra et al., 2020. For each image, the longest βIII tubulin- or Tau-positive neurite was selected and straightened (*File* > *Open* [*do not autoscale*]; *Segmented Line*; *Selection* > *Straighten*). An in-house FIJI/ImageJ macro (*Concentric_circles*) was used to generate concentric circles at 10 μm intervals from the soma edge covering up to 150 μm of the neurite whenever possible. Fluorescence intensity was measured in each bin. Additionally, background intensity was measured and subtracted.

For puromycin intensity profiles at EV-axon contact points, a Tau-positive axon was selected in each relevant channel (PKH26 and puromycin) using the Segmented Line tool. The fluorescence profile was generated (*Analyze* > *Plot Profile*), setting the 0 μm point at the pixel with the highest PKH26 intensity. Puromycin intensity was measured within 5 μm in both directions. For control regions far from EV-axon contact sites, the 0 μm point was randomly assigned.

#### 16.3. Fluorescence discrete puncta analysis in binarized images

Pre- and postsynaptic compartments were identified with anti-synaptophysin and anti-Homer-1 antibodies respectively, and quantified in FIJI/ImageJ. βIII tubulin-positive neurites with an average length of 50 – 70 μm were selected and straightened (*Segmented Line*; *Selection* > *Straighten*). Each neurite was then binarized (*Image* > *Adjust* > *Threshold* > *MaxEntropy; Process* > *Binary* > *Make Binary*) and puncta were quantified using the Analyze tool (Size: 0.05 – 1 μm) ^31^.

### 17. Statistical Analyses

Statistical analyses were performed using Prism 8 or Prism 10 software (GraphPad Software, Dotmatics. MA, USA). Whenever possible, a randomized block design was followed to control inter-experimental variability. Comparisons between two groups were conducted using a two-tailed t-test, while one- or two-way ANOVA was used for multiple group comparisons. If an outlier value was suspected, data were subjected to the Grubb’s test (α=0.05). Sample size and statistical details are provided in the figure legends. For all cases, a p-value < 0.05 was considered statistically significant.

## Results

### 1. Astrocytes modulate local protein synthesis in neurons

To determine if astrocytes regulate local translation in neurons, we first addressed whether the presence of astroglia in neuronal cultures affected the synthesis of neuritic proteins. We performed primary neuronal cultures in modified Boyden chambers in the absence or presence of primary astrocytes. Boyden chambers enable the characterization of molecular components (e.g. proteins, RNAs) present in distal neurites (lower compartment in **Figure 1A^i^**) separately from the molecular content of proximal projections, somata and nuclei (upper compartment in **Figure A^i^**) ^32^. Indeed, our own immunoblotting analyses confirmed the enrichment of the neuronal specific marker βIII tubulin in distal neurites (Neurite in **Figure 1A^ii^**), while the transcription factor NeuN was abundantly found in proximal neuronal compartments (Soma in **Figure 1A^i^**) but not in neuritic extracts. These results were consistent with the absence of nuclei, detected by immunofluorescence, in the lower chamber compartment (Neurite in **Figure 1A^iii^**). Finally, most of distal neurites were identified as axons based on Tau immunoreactivity, while dendrites (positive for Map2) were found to a lesser extent (**Figure 1A^iii^**). Once we had validated in our own hands the isolation of subneuronal compartments in Boyden chambers, we analyzed the levels of newly synthesized neuritic proteins in neurons cultured in the absence (culture setup 1, **Figure 1B^i^**) or presence of astrocytes (culture setup 2, **Figure 1B^i^**). It is worth noting that in this culture system, neurons and glia communicate with each other through their secretome, as no direct contact between cell types is established (see Methods section). On another side, evidence shows that amyloid pathology, which is central to the development of Alzheimer’s disease, enhances local protein synthesis in axons ^16^. Thus, we used amyloid-β peptides (oligomeric Aβ_1-42_) to treat both neuronal monocultures and neuron-astrocyte co-cultures for 24 hours as a positive control (**Figure 1B**). Cells were labelled with O-propargyl-puromycin (OPP) to detect newly synthesized proteins. After protein extraction, OPP was conjugated with biotin and proteins were processed for immunoblotting. OPP-biotin conjugates in neuritic extracts were revealed with streptavidin. Results indicate that while no significant protein synthesis was observed in neurites from monocultures treated with vehicle (DMSO) compared to non-OPP treated cells, OPP-biotin labelled polypetides were readily detectable in control neurons cultured in the presence of astrocytes, as well as in monocultures and co-cultures exposed to Aβ (**Figure 1B^ii^**, left graph). Additionally, a significant increase in newly synthesized neuritic proteins was observed in neuron-astrocyte co-cultures compared to neuronal monocultures in basal conditions, while no changes were detected between both cultures in the presence of Aβ (**Figure 1B^ii^**). We then ask if the positive detection of OPP-biotin conjugates was a result of local mRNA translation in neurites. We performed OPP labelling in severed neurites devoid from somatic input to ensure that newly synthesized proteins arose from localized transcripts rather than being transported from the soma. Interestingly, OPP-biotin conjugates were only consistently and unambiguously detected in neurites isolated from neuron-astrocyte co-cultures exposed to Aβ oligomers, and a trend towards an increased was observed in these cultures compared to neuronal monocultures (**Figure 1B^iii^**). These results suggest that astrocytes sustain local protein synthesis in neurons, at least in amyloid induced conditions. In light of these observations, we further focused on glia-mediated regulation of local translation in response to Aβ.

**Figure 1.**
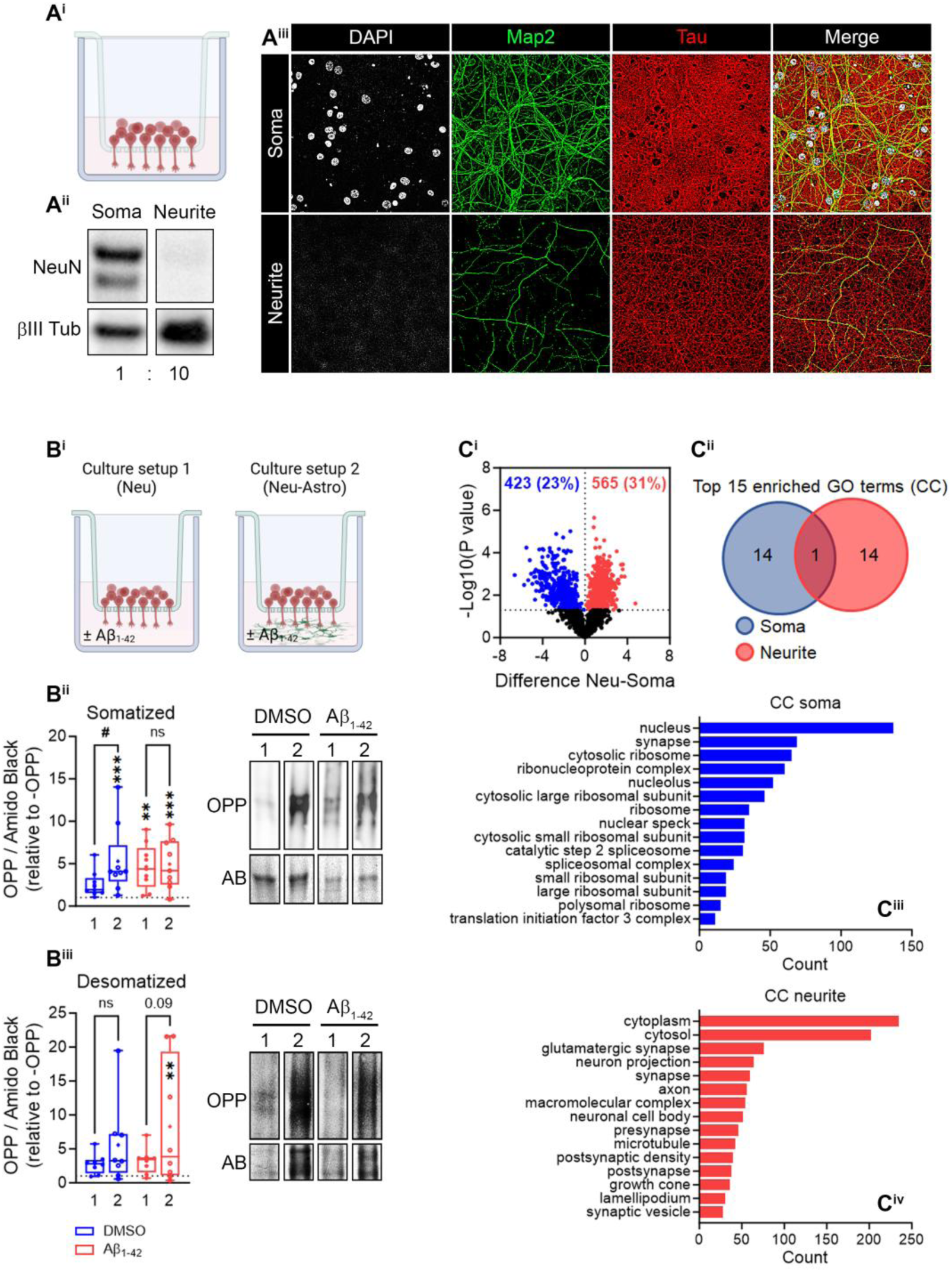
Isolation of neuritic extracts in modified Boyden chambers. **(A)** Boyden chambers enable the separation of distal neurites (lower or neuritic compartment) from their soma, nuclei and proximal dendrites and axons (upper or somatic compartment) Schematic representation of a Boyden chamber created in https://BioRender.com (**A^i^**). The transcription factor NeuN is heavily enriched in the somatic compartment (Soma) but not in distal neurites (Neurite) (**A^ii^**). Nuclei (stained with DAPI) are absent in the neuritic compartment (lower micrographs) of Boyden chambers. Axons (Tau+) are more abundant than dendrites (Map2+) in the neuritic compartment (lower micrographs) (**A^iii^**). **(B)** Detection of newly synthesized proteins in neuritic extracts by O-propargyl puromycin (OPP) labelling. Schematic representation of neuronal cultures (left) and neuron-astrocyte co-cultures (right) in Boyden chambers. Created in https://BioRender.com (**B^i^**). The box and whisker graph (left) represents the levels of OPP measured by streptavidin in neurites form somatized cells in 9 independent cultures (N=9). Data are normalized to total protein stained with Amido Black and are represented relative to the -OPP condition (dashed line). 1, neuronal cultures; 2, neuron-astrocyte co-cultures. One-way ANOVA followed by Holm-Šídák *post hoc* test for selected pairs of columns. Asterisks represent significant changes detected between each experimental condition and the -OPP negative control: **p < 0.01; ***p < 0.001; n.s, not significant. The hash sign indicates significant changes detected in neuron-astrocyte co-cultures (2) compared to monocultures (1) in control or Aβ-treated cells: #p < 0.05; n.s, not significant. Representative images of western blotting membranes stained for streptavidin and Amido Black (AB) are shown (right panel) (**B^ii^**). The box and whisker graph (left) represents the levels of OPP measured by streptavidin in neurites form desomatized cells in 8 independent cultures (N=8). Data are normalized to total protein stained with Amido Black and are represented relative to the -OPP condition (dashed line). 1, neuronal cultures; 2, neuron-astrocyte co-cultures. One-way ANOVA followed by Holm-Šídák *post hoc* test for selected pairs of columns. Asterisks represent significant changes detected between each experimental condition and the -OPP negative control: *p < 0.05; n.s, not significant. OPP levels in neuron-astrocyte co-cultures (2) compared to monocultures (1) in control or Aβ-treated cells are also indicated. Representative images of western blotting membranes stained for streptavidin and Amido Black (AB) are shown (right panel) (**B^iii^**). **(C)** Protein enrichment in neuritic and somatic extracts. The volcano plot represents the upregulation (red dots) and downregulation (blue dots) of proteins detected in distal neurites compared to the somatic compartment. Results are the average of 4 independent experiments (N=4) (**C^i^**). Venn diagram depicting the overlap of proteins clusters grouped with DAVID software based on cellular components (CC) (**C^ii^**). Top 15 enriched protein clusters in the soma (**C^iii^**) and in neurites (**C^iv^**) based on the CC (p < 0.05).

### 2. Astrocytes modulate the levels of ribosomal proteins localized to distal neurites

A pre-requisite for local protein synthesis in neurons is the transport of both mRNAs and components of the translation machinery to distal neurites (e.g. dendrites, axons). Thus, we performed proteomic analyses to determine if translation regulators were present in neurites, and whose levels could be influenced by glia. We performed primary neuronal cultures in modified Boyden chambers and neuritic proteins were isolated separately from those in the somatic compartment (which includes neuronal somata, nuclei and proximal neurites) and subjected to mass spectrometry. We first analyzed the identity of proteins enriched in both extracts (regardless of the presence or absence of astrocytes and of the treatment with vehicle or Aβ) and observed that 31% (565) of them were significantly increased in neurites compared to the somatic compartment, whereas 23% (423) were decreased (**Figure 1C^i^**). We then clustered the significantly changed proteins into GO terms based on the cellular component (CC). Only one cluster was commonly found in both compartments (**Figure 1C^ii^**) indicating the distinct nature of proteins selectively localized to neurites. We further analyzed those GO terms and observed that proteins enriched in the somatic compartment sustained nuclear functions (including that of the nucleolus, the nuclear speck and the spliceosome), cytoplasmic functions (including cytoplasmic translation) and synapses (**Figure 1C^iii^**). Conversely, proteins enriched in neurites were mainly related to neuronal projections (including axons, growth cones and lamellipodia) and synapses (both pre- and postsynapses) (**Figure 1C^iv^**). These results confirmed that the selective separation of subneuronal fractions in Boyden chambers is suitable to assess the regulation of neuritic proteins by mass spectrometry.

We then asked if the presence of astrocytes in neuronal cultures modulated the local neuritic proteome. We performed proteomic analyses from neuritic extracts isolated from monocultures and neuron-astrocyte co-cultures and took advantage of another unrelated dataset in which we aimed to address the influence of microglia on the local proteome. Both datasets contained information on the up- or downregulation of neuritic proteins compared to the somatic compartment in neuronal monocultures. We focused on those which showed a similar trend in both datasets (red and blue dots in **Supplementary Figure 1A**) and discarded proteins whose relative levels in neurites compared to the soma showed opposite trends in both sets of experiments (black dots in **Supplementary Figure 1A**). After filtering, we analyzed the identity of neuritic proteins affected by both astrocytes and microglia or by either glial cell type. Hence, we focused on proteins significantly changed in neuron-glia co-cultures compared to neuronal monocultures. We observed an overlap of 500 proteins regulated by both glial cell types, whereas 234 were exclusively regulated by astrocytes and 631 by microglia (**Supplementary Figure 1B**). Among the commonly regulated proteins, most of them were involved in translation, protein transport or vesicle transport (endo- and/or exocytosis) (grey bar graph, **Supplementary Figure 1Ci**). On the other hand, astrocytes specifically affected protein biogenesis (e.g. translation, protein folding, ribosome assembly…) (green bar graph, **Supplementary Figure 1Cii**), whereas microglia were mostly involved in transport (e.g. protein transport, vesicular transport…) and organization of neuronal projections (e.g. axonogénesis, cell polarity, cytoskeleton organization…) (purple bar graph, **Supplemantary Figure 1C^iii^**). Our results thus suggest that astrocytes and microglia sustain common biological processes (BP) but also selectively regulate distinct protein cohorts in neurites, with astroglia being mainly involved in protein synthesis-related functions.

We then selected proteins included in all significantly enriched biological functions (BPs) regulated by astrocytes (cyan bar graph, **Supplementary Figure 1Cii**) and performed protein-protein interaction analyses with STRING. Interestingly, the most represented functional network was that of translation regulation (MCL=1.4), which included ribosomal proteins and translation initiation factors (**Figure 2A**). Most of the proteins within this cluster were upregulated in neurites isolated from neuron-astrocyte co-cultures compared to neuronal monocultures, especially in response to Aβ oligomers (**Figure 2B^i^**). For instance, according to proteomic analyses, Rpl26, a component of the S60 ribosome located at the surface of the large subunit, was significantly increased in the presence of astrocytes both in vehicle and Aβ-treated cultures (**Figure 2B^ii^**, left graph). Conversely, Rps6, a component of the S40 subunit was upregulated only in Aβ-treated co-cultures and not in basal conditions (**Figure 2B^ii^**, right graph). It is worth noting that the presence of Rps6 in axons at its phosphorylation has been historically used to address local translation activation. Changes in Rpl26 were partially validated by immunocytochemistry analyses (**Figure 2C^i^**), in which we observed that protein levels were indeed enhanced in neuro-astrocytes co-cultures in distal axons (ca. 150 µm away from the soma. **Figure 2C^i^** and inset in **C^ii^**), although only upon Aβ treatments and not in control conditions. However, Rps6 increased levels in distal axons (ca. 150 µm away from the soma; insets in **Figure 2D^ii^**) measured by immunostaining were consistent with the proteomic data in which the effect was only observed in response to Aβ (right graph in **Figure 2B^ii^**). These results, together with the ones reported in the previous epigraph, suggest that neuron-astrocyte communication through secreted factors modulate local translation in neurons.

**Figure 2.**
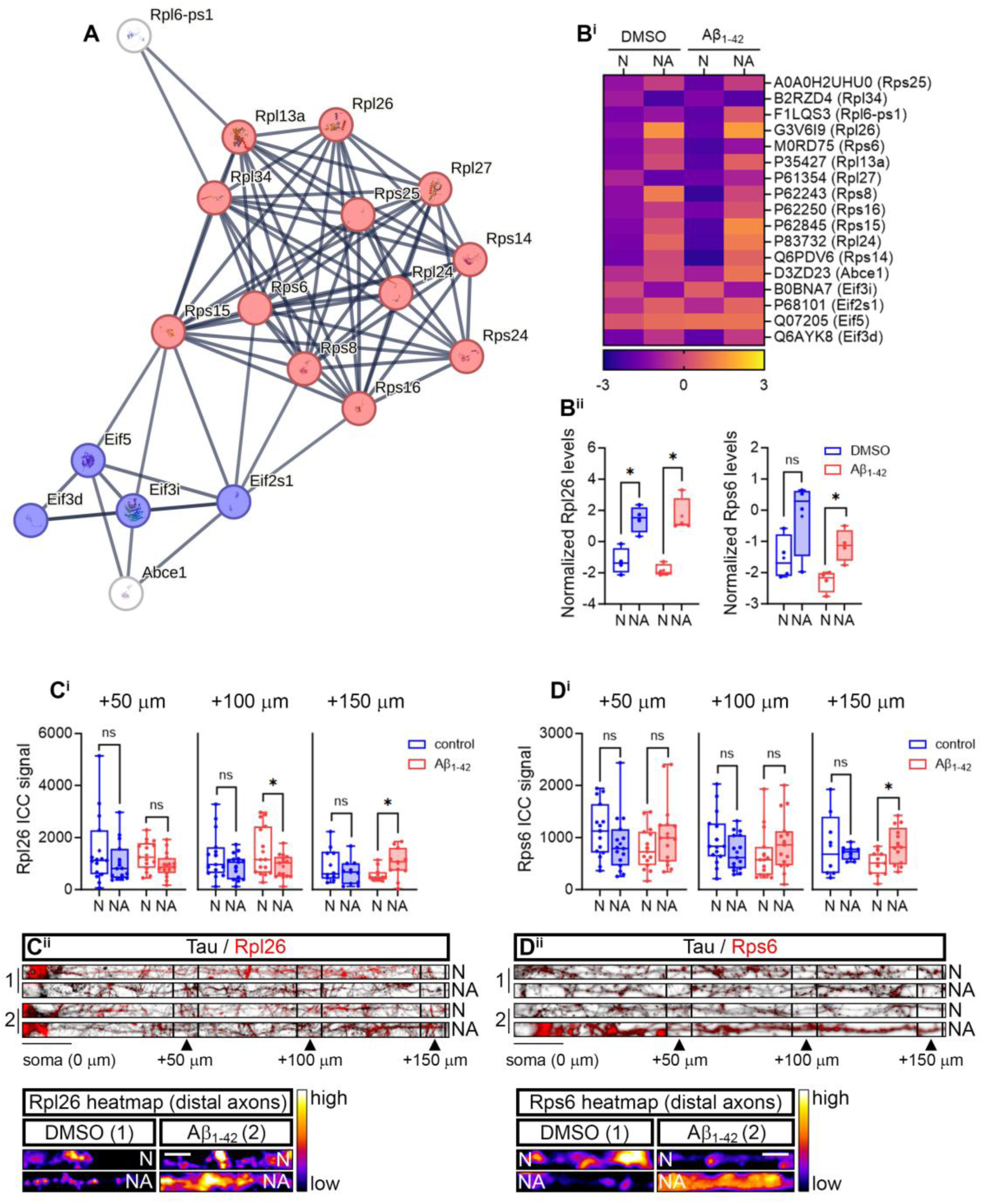
The presence of astrocytes in culture significantly affects levels of translation regulators in distal neurites. **(A)** STRING diagram representing the largest functional network (MCL=1.4) changed by the presence of astrocytes in culture (regardless of the treatment with vehicle or Aβ oligomers). **(B)** Normalized levels of translation regulators in neuritic extracts. The heatmap shows the levels of significantly regulated proteins (depicted in **(A)**) in co-cultures (NA) compared to monocultures (N) in control and Aβ-treated cells. The graph represents the average of 4 independent experiments (N=4) (**B^i^**). Box and whisker graphs indicate the normalized levels of ribosomal proteins Rpl26 (left) and Rps6 (right) obtained by proteomics from 4 independent cultures (N=4). Pairwise t-tests comparing co-cultures (NA) to monocultures (N) in control or Aβ-treated cells. *p < 0.05; n.s, not significant (**B^ii^**). **(C)** Evaluation of Rpl26 in axons by immunocytochemistry. Box and whiskers graphs show Rpl26 fluorescent levels in axons from neurons cultured alone (N) or in the presence of astrocytes (NA) upon vehicle or Aβ exposure. Results represent the fluorescent signal of 10-15 (n=10-15) axonal segments at different distances from the soma measured in three independent cultures (N=3). Pairwise t-tests comparing co-cultures (NA) to monocultures (N) in control or Aβ-treated cells. *p < 0.05; n.s, not significant (**C^i^**). Micrographs representing the results obtained in (**C^i^**) are shown (**C^ii^**). Insets in (**C^ii^**) show heatmaps from Rpl26 fluorescent signal in distal axons (ca. 150 µm from the soma). Scale bar = 2µm. **(D)** Evaluation of Rps6 in axons by immunocytochemistry. Box and whiskers graphs show Rps6 fluorescent levels in axons from neurons cultured alone (N) or in the presence of astrocytes (NA) upon vehicle or Aβ exposure. Results represent the fluorescent signal of 10-15 (n=8-14) axonal segments at different distances from the soma measured in three independent cultures (N=3). Pairwise t-tests comparing co-cultures (NA) to monocultures (N) in control or Aβ-treated cells. *p < 0.05; n.s, not significant (**D^i^**). Micrographs representing the results obtained in (**D^i^**) are shown (**D^ii^**). Insets in (**D^ii^**) show heatmaps from Rpl26 fluorescent signal in distal axons (ca. 150 µm from the soma). Scale bar = 2µm.

### 3. Effect of astroglial EVs on the *de novo* synthesis of neuritic proteins

Given that our results thus far indicate that the secretome released in neuron-astrocyte co-cultures regulate local protein synthesis in neurons, we wondered if this effect could be driven, at least partially, by astroglial EVs secreted in basal conditions and/or in response to Aβ peptides. We cultured primary astrocytes in serum-free medium (3 DIV, as in co-cultures) and treated them with vehicle or Aβ oligomers for 24 hours. Given the limited amount of sample, we collected all EVs secreted by the cells by using a single ultracentrifugation at 250,000 x g from the conditioned media and the presence of canonical EV markers was analyzed by immunoblotting. For comparative controls we also characterized neuronal EVs. CD63, TSG101 and Hsp70 were present in vesicular extracts from control- and Aβ-treated astroglial cultures (**Figure 3A**, 2-most-left panels), whereas the endoplasmic reticulum-associated protein Grp78 (**Figure 3A**, 2-most-left panels) was detected in this fraction. Similar results were obtained for neuronal EVs (**Figure 3A**, 2-most-right panels). We then addressed the size and shape of astroglial and neuronal EVs by cryo-EM. Most particles released by astrocytes and neurons (regardless of the treatment with Aβ) were smaller than 200 nm-diameter in size but larger than 150 nm (**Figure 3B^ii^**). We classified these nanoparticles into four groups based on their shape and the electrodensity of their membrane (**Figure 3B^i^**): type I vesicles were spherical, with high-electrodensity membranes often “decorated” at their outer surface; type II vesicles showed high-electrodensity membranes enclosed within a larger membrane, resembling multivesicular bodies; type III vesicles were classified as irregular, non-spherical particles with lower-electrodensity membranes, which reminded of autophagolysomes; and type IV particles showed a disrupted membrane and were considered apoptotic bodies (based on their larger size) or as EVs that were disturbed during the ultracentrifugation step. These cryo-EM images provide evidence that most particles released by cells to the extracellular medium are predominantly composed of vesicular structures, indicating a high content of EVs. Interestingly, almost 80% of astroglial vesicular extracts contained type I particles, whereas neuronal EVs were more heterogeneous in shape (**Figure 3B^iii^**). Finally, we performed nanoparticle tracking analyses (NTA) to address the effect of Aβ on EV release. Size-range comparison of EVs released by different cell types confirmed that neuronal EVs were more heterogeneous than astroglial EVs (data not shown). We then observed that, in line with the cryo-EM results, most astroglial EVs had a smaller diameter than 200 nm. Aβ oligomers enhanced the overall release of EVs from astrocytes as reported by both NTA (**Figure 3C^i^**) and the area under the curve (AUC, inset in **Figure 3C^i^**). Aβ did not affect the average size of astroglial EVs (**Figure 3C^ii^**) but a significant increase was observed in the secretion of the major EV population (right graph in **Figure 3C^iii^**; peak size of NTA curves). Similarly, overall neuronal EV secretion was significantly increased by Aβ peptides (**Figure 3D^i^**), but no differences were observed neither in the average size (**Figure 3D^ii^**) nor in the release of EVs at the peak size of NTA curves (**Figure 3D^iii^**), although a trend towards an increase was observed in the latter parameter.

**Figure 3.**
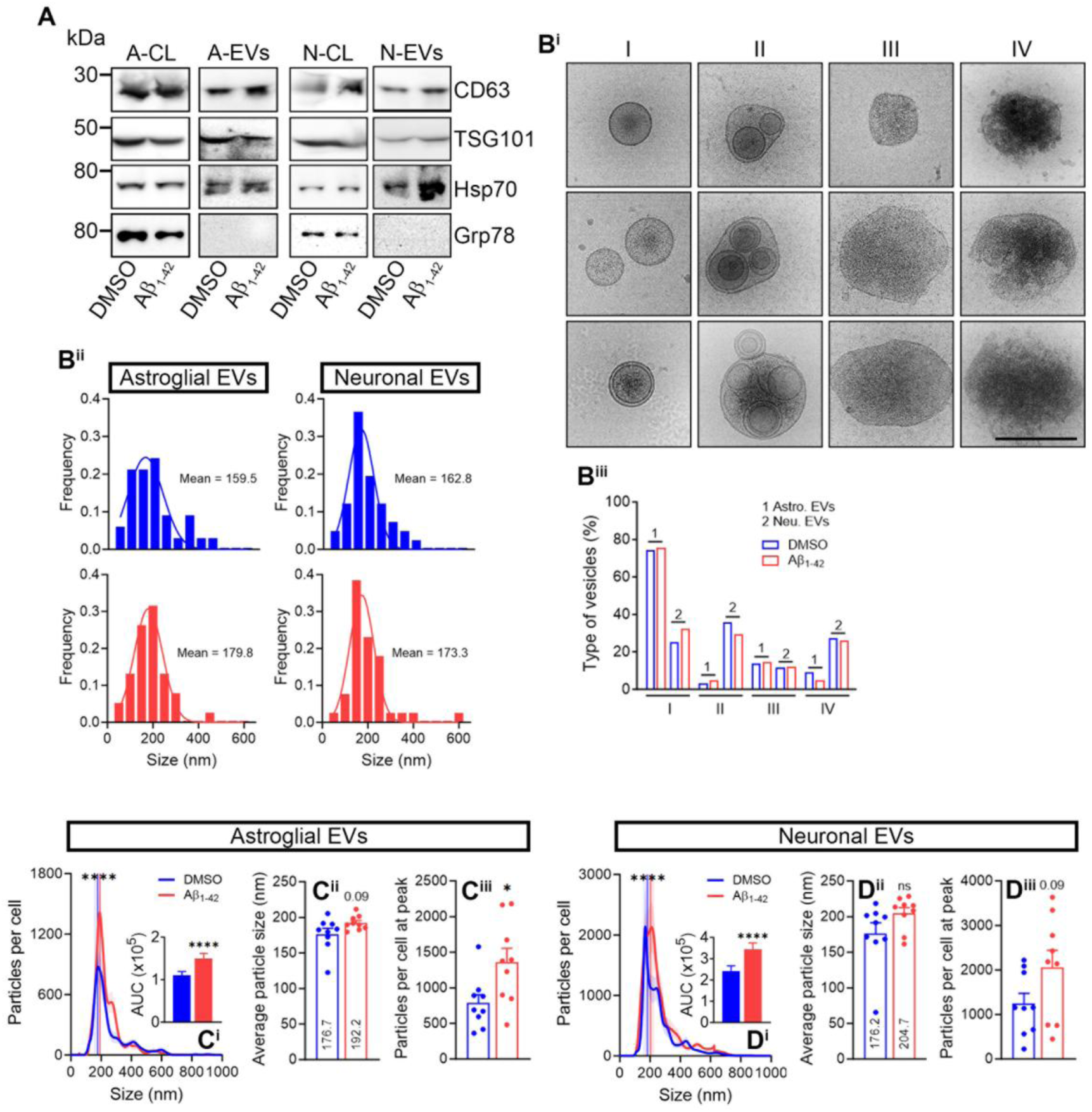
Characterization of EVs released by astrocytes and neurons in control conditions and in response to Aβ oligomers. **(A)** Western blots showing the presence of EV markers in astroglial and neuronal whole cell lysates (A-CL and N-CL respectively) and in astroglial and neuronal vesicular extracts (A-EVs and N-EVs respectively). Note that the rough endoplasmic reticulum protein Grp78 was only detected in whole lysates but not in EVs. **(B)** Visualization of astroglial and neuronal EVs by Cryo-EM. EVs were classified based on their size, shape and electrodensity. Type I vesicles are described as < 200 nm in diameter spherical particles, with highly electrodense single or double membranes often “decorated” at their outer surface. Type II vesicles showed highly electrodense membranous particles enclosed within a larger membrane, resembling multivesicular bodies. Type III vesicles were classified as irregular, non-spherical particles with lower electrodense membranes. Type IV particles, typically larger than 200 nm in diameter, showed a disrupted membrane. Scale bar = 200 nm (**B^i^**). Qualitative assessment of size distribution of atroglial and neuronal EVs released in control conditions (blue frequency graphs) and in response to Aβ (red frequency graphs). Graphs specify the average size of EVs released in each experimental condition (**B^ii^**). Graph representing the percentage of different types of EVs released by astrocytes and neurons in control conditions and upon Aβ treatment (**B^iii^**). **(C)** Quantification of EVs released by astrocytes by nanoparticle tracking analysis (NTA). NTA curves are normalized to the number of secreting cells. Results were analyzed by Two-way ANOVA. ****p < 0.0001 (**C^i^**). The area under the curve (AUC) graph retrieved from NTA data shows overall increased particle secretion by astrocytes exposed to Aβ peptides. t-test analysis; ****p < 0.0001 (inset in (**C^i^**)). The bar graph represents the average size ± SEM of EVs released by control and Aβ-treated astrocytes measured in 9 vesicular preparations (n=9) isolated from 3 independent experiments (N=3). t-test analysis; n.s, not significant (**C^ii^**). A bar graph representing the number of particles ± SEM belonging to the major EV population (particles per cell at peak of the NTA curve) is shown. t-tests were performed in in 9 vesicular preparations (n=9) isolated from 3 independent experiments (N=3). *p < 0.05 (**C^iii^**). **(D)** Quantification of EVs released by neurons by nanoparticle tracking analysis (NTA). NTA curves are normalized to the number of secreting cells. Results were analyzed by Two-way ANOVA. ****p < 0.0001 (**D^i^**). The area under the curve (AUC) graph retrieved from NTA data shows overall increased particle secretion by neurons exposed to Aβ peptides. t-test analysis; ****p < 0.0001 (inset in (**D^i^**)). The bar graph represents the average size ± SEM of EVs released by control and Aβ-treated neurons measured in 9 vesicular preparations (n=9) isolated from 3 independent experiments (N=3). t-test analysis; n.s, not significant (**D^ii^**). A bar graph representing the number of particles ± SEM belonging to the major EV population (particles per cell at peak of the NTA curve) is shown. t-tests were performed in in 9 vesicular preparations (n=9) isolated from 3 independent experiments (N=3). The p value obtained in the statistical comparison is shown (**D^iii^**).

We then treated primary neurons with EVs released by control- (hereafter DMSO-EVs) or Aβ-treated (from now on Aβ-EVs) astrocytes for 45 minutes and measured puromycin labelling along axons. Treatment with fresh medium devoid of EVs (FM; -EVs) was used to establish baseline levels on newly synthesized proteins. Our results indicate that overall puromycin intensity in axons was increased by Aβ-EVs but not by DMSO-EVs (**Figure 4A^i^**, left graph and **Figure 4A^ii^**). We then focused on specific distances from the neuronal soma. Results revealed that Aβ-EVs enhanced puromycilated proteins in distal axons, yet DMSO-EVs decreased their levels, (ca. 130-150 µm from the soma; **Figure 4A^i^**, right graph and insets in **Figure 4A^ii^**), suggesting a distinct local effect of astroglial EVs depending on the experimental context they were released. We used the same approach with neuronal EVs for comparative purposes. Surprisingly in this case, general puromycin levels were enhanced by DMSO-EVs but not by Aβ-EVs (**Figure 4B^i^**, left graph and **Figure 4B^ii^**), and the former seemed to selectively affect newly synthesized proteins in distal axons (ca. 150 µm from the soma; **Figure 4B^i^**, right graph and insets in **Figure 4B^ii^**). Thus far, our results suggest that not only the context but also the cell types releasing EVs differentially modulate local translation in axons.

**Figure 4.**
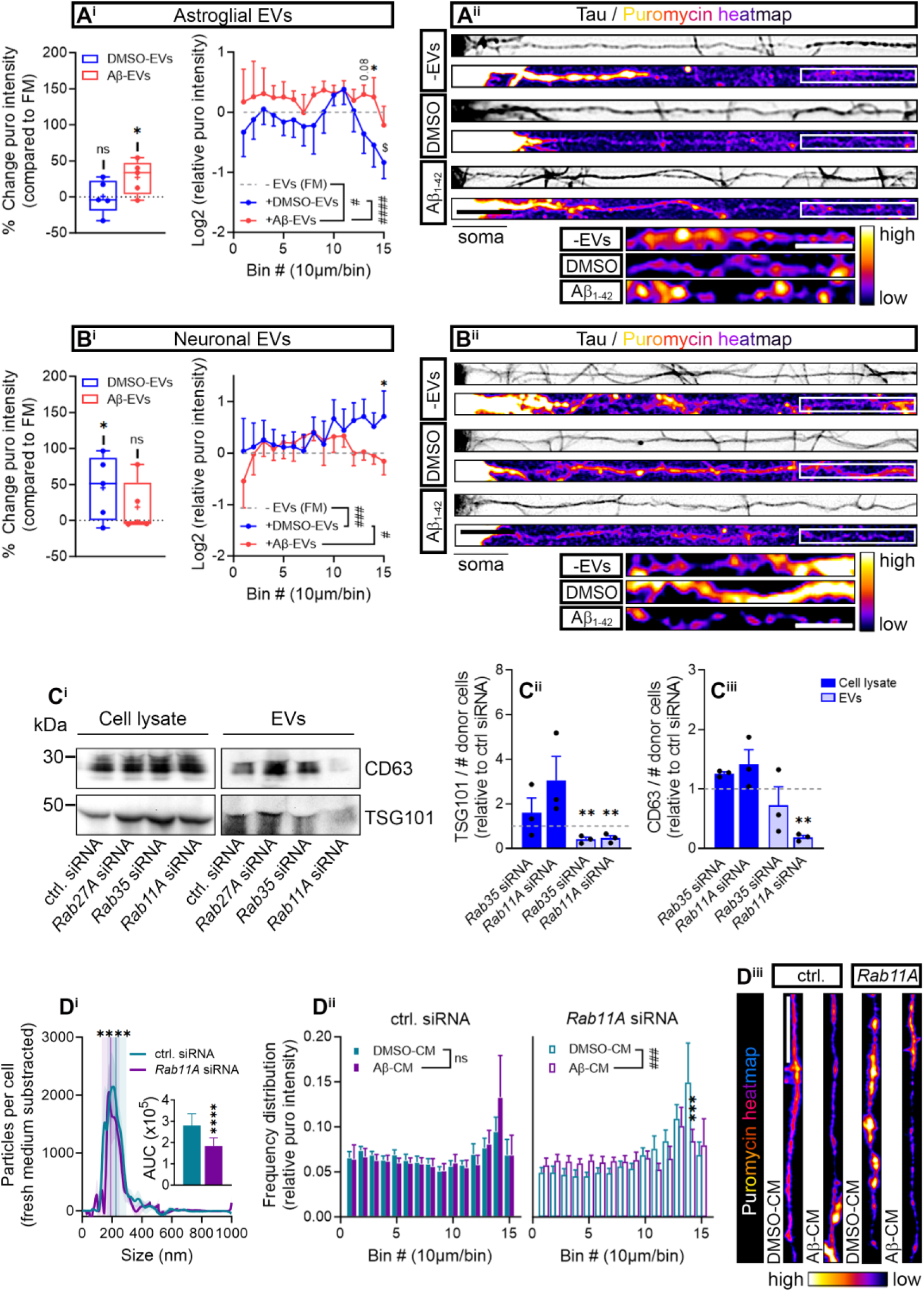
Effect of EVs released by astrocytes in newly synthesized axonal proteins. **(A)** EVs released by control and Aβ-treated astrocytes (DMSO-EVs and Aβ-EVs respectively) differentially affect puromycin levels in axons. Box and whisker graph representing percent changes in puromycin fluorescent intensity in astroglial DMSO-EV- and Aβ-EV-treated neurons compared to fresh medium-treated neurons (FM). One-way ANOVA followed by Dunnet’s *post hoc* test comparing the effect of DMSO- or Aβ-EVs to FM. *p < 0.05; n.s, not significant (**A^i^**; left graph). Intensity vs distance graph representing puromycin intensity in axons measured in 15 10-µm bins covering a distance of 150 μm from the soma. Data are plotted as puromycin intensity relative to the FM condition (Log2 was applied for better visualization in the graph). Two-way ANOVA followed by Holm-Šídák *post hoc* test. The hash signs indicate overall changes detected in all experimental groups. #p < 0.05; ####p < 0.0001. Asterisks depict changes in DMSO- or Aβ-EV-treated neurons compared the the FM. *p < 0.05. The dollar sign indicates changes in neurons treated with DMSO- and Aβ-EVs. $p < 0.05. Statistical analyses were performed in results retrieved from 5 independent experiments (N=5) (**A^i^**; right graph). Micrographs representing the results obtained in (**A^i^**) are shown (**A^ii^**). Insets in (**A^ii^**) show heatmaps from puromycin fluorescent signal in distal axons. Scale bar = 20 µm (10 µm in insets). **(B)** EVs released by control and Aβ-treated neurons (DMSO-EVs and Aβ-EVs respectively) differentially affect puromycin levels in axons. Box and whisker graph representing percent changes in puromycin fluorescent intensity in astroglial DMSO-EV- and Aβ-EV-treated neurons compared to fresh medium-treated neurons (FM). One-way ANOVA followed by Dunnet’s *post hoc* test comparing the effect of DMSO- or Aβ-EVs to FM. *p < 0.05; n.s, not significant (**B^i^**; left graph). Intensity vs distance graph representing puromycin intensity in axons measured in 15 10-µm bins covering a distance of 150 μm from the soma. Data are plotted as puromycin intensity relative to the FM condition (Log2 was applied for better visualization in the graph). Two-way ANOVA followed by Holm-Šídák *post hoc* test. The hash signs indicate overall changes detected in all experimental groups. #p < 0.05; ###p < 0.001. Asterisks depict changes in DMSO- or Aβ-EV-treated neurons compared the the FM. *p < 0.05. Statistical analyses were performed in results retrieved from 5 independent experiments (N=5) (**B^i^**; right graph). Micrographs representing the results obtained in (**B^i^**) are shown (**B^ii^**). Insets in (**B^ii^**) show heatmaps of puromycin fluorescent signal in distal axons. Scale bar = 20 µm (10 µm in insets). **(C)** *Rab11A* genetic downregulation in astrocytes reduces vesicular markers in EVs. Western blots depicting the levels of CD63 and TSG101 (normalized to the number of secreting astrocytes) in cell lysates and EVs released upon transfection of astroglial cultures with control, *Rab27A*, *Rab35* or *Rab11A* siRNAs (**C^i^**). Bar graphs represent the levels relative to control-transfected cells (dashed lines) of TSG101 (**C^ii^**) and CD63 (**C^iii^**) in cell lysates and EVs, upon downregulation of Rab35 and Rab11A in astrocytes, measured in three independent experiments (N=3). One-way ANOVA (performed separately in cell lysates and EVs) followed by Dunnet’s *post hoc* comparison (*Rab35* or *Rab11A* siRNAs vs control siRNA). **p < 0.01. **(D)** Rab11A inhibition alters EV release by astrocytes and disturbs EV-dependent puromycin distribution in neuronal axons. Quantification of EVs released by astrocytes by NTA. Data from fresh medium treated with lipofectamine were subtracted from the curves and resulting values were normalized to the number of secreting cells. Two-way ANOVA. ****p < 0.0001 (**D^i^**). The area under the curve (AUC) graph retrieved from NTA data shows overall decrease in particles released by astrocytes transfected with *Rab11A* siRNA. t-test analysis; ****p < 0.0001 (inset in (**D^i^**)). Intensity vs distance graph representing puromycin frequency distribution in axons measured in 15 10-µm bins covering a distance of 150 μm from the soma in control. Neurons were treated with conditioned media from astrocytes transfected with control siRNA (left) or *Rab11A* siRNA. Data corresponds to 6 independent experiments (N=6). Two-way ANOVA followed by Holm-Šídák *post hoc* test. The hash sign indicates overall changes between axons treated with conditioned media from control and Aβ-treated astrocytes: ###p < 0.001; n.s, not significant. Asterisks indicate significant changes at a given distance from the soma: ***p < 0.001 (**D^ii^**). Micrographs representing the results quantified in (**D^ii^**) are shown. Scale bar = 20 µm (**D^iii^**).

To further confirm that astroglial Aβ-EVs indeed positively regulated the synthesis of proteins targeted to distal axons, we genetically inhibited molecules involved in EV biogenesis and evaluated the effect of astrocyte-derived conditioned medium (CM) on neurons. Primary astrocytes were transfected with a control siRNA or with siRNAs targeting *Rab27A, Rab35* or *Rab11A*. Although no efficient knockdown was observed in astrocytes treated with *Rab27A* siRNA, *Rab35* and *Rab11A* siRNAs significantly decreased their corresponding targets at the mRNA (**Supplementary Figure 2A**) and protein levels (**Supplementary Figure 2B**). Indeed, *Rab27A* siRNA did not seem to affect vesicular markers either (**Figure 4C^i^**), thus we focused on the inhibition of Rab35 and Rab11A. Decreased levels of TSG101 in vesicular extracts were detected upon both *Rab35* and *Rab11A* knockdown (**Figure 4C^ii^**), however CD63 was only reduced in EVs released by *Rab11A* siRNA-treated astrocytes (**Figure 4C^iii^**). Given the more robust effect of *Rab11A* siRNA in astroglial EV secretion, we further focused on this protein in following experiments. NTA analyses revealed significant changes in the size distribution of EVs released by *Rab11A* siRNA-transfected astrocytes compared to control transfections (**Figure 4D^i^**) and decrease in their secretion (AUC; inset in **Figure 4D^i^**). Finally, we treated neurons either with conditioned media from untransfected astrocytes or with conditioned media from astroglial cultures transfected with control or *Rab11A* siRNAs. In all cases astrocytes were treated with vehicle or Aβ. We observed that neither conditioned media secreted by unstransfected cells changed overall puromycin levels in axons compared to fresh medium (FM; **Supplementary Figure 3Ai**). Additionally, no differences in the relative puromycin distribution along axons was observed when neurons were treated with conditioned medium from vehicle- or Aβ-treated astrocytes (**Supplementary Figure 3Aii**). Similar results were observed in the relative puromycin distribution when conditioned media were collected from control-transfected astrocytes (left graph in **Figure 4D^ii^** and **Figure 4D^iii^**). In contrast, a significant decrease in puromycin was observed in distal axons (ca. 140 µm from the soma) when neurons were treated with conditioned medium from Aβ-treated astrocytes transfected with *Rab11A* siRNA (right graph in **Figure 4D^ii^** and **Figure 4D^iii^**). All in all, these results confirm that astroglial Aβ-EVs released in a Rab11A-dependent manner enhance de *novo synthesis* of proteins targeted to distal axons.

Experiments performed thus far aimed at addressing the direct effect of astroglial Aβ-EVs on protein synthesis in axons, hence the 45-minute treatments. However, we then wondered if glial EVs could elicit a more long-lasting effect on neurons. To answer this question, we treated neuronal cultures with EVs for 24 hours and observed a significant increase in axonal puromycin levels when cells were exposed to Aβ-EVs compared to DMSO-EVs (**Supplementary Figure 3B**, left graph). Interestingly, no differences were detected when cells were treated with neuronal DMSO-EVs or Aβ-EVs (**Supplementary Figure 3B**, right graph). We then used a complementary approach to confirm the long-term effect of astroglial Aβ-EVs on newly synthesized proteins. Namely, neurons were treated with conditioned media depleted from DMSO-EVs or Aβ-EVs. When astroglial medium was depleted from DMSO-EVs a significant increase in protein synthesis was observed compared to the complete medium, while depletion of Aβ-EVs led to a decrease (left graph in **Supplementary Figure 3C** and fluorescence micrographs), however in this case EV-depletion did not affect only distal but also proximal sites of axons (right graph in **Supplementary Figure 3C** and fluorescence micrographs). Conversely no changes in protein synthesis were detected when neuronal media was depleted from DMSO-EVs or Aβ-EVs (**Supplementary Figure 3D**). These results confirm that the positive regulation of newly synthesized axonal proteins by astroglial Aβ-EVs is long-lasting.

### 4. Astroglial Aβ-EVs enhance local protein synthesis in axons and upregulate synaptic markers

The facts that 1) desomatized neurites were able to locally incorporate puromycin in Aβ-treated neuron-astrocytes co-cultures (**Figure 1B^ii^**), 2) ribosomal proteins Rpl26 and Rps6 were found increased in distal axons from Aβ-treated neuron-astrocytes co-cultures (compared to neuronal monocultures; **Figures 2B^ii^-D^ii^**), and 3) puromycin signal showed increased staining ^33,34^ in distal axons exposed to astroglial Aβ-EVs (**Figure 4A**), all strongly indicate that glial vesicles released in response to Aβ oligomers enhance local translation in neurons. However, to unambiguously demonstrate the local effect of astrocyte-derived EVs and to rule out the potential transport of somatically synthesized proteins targeted to the neuronal periphery, we cultured primary neurons in microfluidic chambers and specifically targeted axons with EVs.

Microfluidic chambers enable the fluidic isolation of axons from their corresponding somatodendritic compartment (**Figures 5A^i^** and **5A^ii^**). Once isolated, axons can be locally treated (e.g. with EVs, puromycin, etc…) without affecting the soma or the dendrites. These microfluidic devices have been used in the past to demonstrate intra-axonal protein synthesis dysregulation in Alzheimer’s disease (AD) ^16,35^ and amyotrophic lateral sclerosis (ALS) ^36^ models. We first analyzed the ability of astrocytes to transfer EVs to axons. To that end, we cultured neurons in microfluidics and once the axons reached the distal compartments, CD63-GFP tagged astrocytes were cultured together with axons. We detected astrocytic-derived CD63-positive particles docked at/or internalized within axons (**Supplementary Figure 4A**). We then treated isolated axons grown in microfluidic chambers with DMSO-EVs or Aβ-EVs released by astrocytes. EVs were labelled with PKH26 lyophilic dye. Qualitative observations revealed a higher PKH26 signal in axonal compartments treated with Aβ-EVs compared to DMSO-EVs. These results are consistent with the NTA quantifications in which we observed a significant increase in EV release by donor cells exposed to Aβ oligomers (**Figure 3C**). However, after a treatment as short as 45 minutes, we did not detect any changes in the docking and/or internalization of DMSO-EVs compared to Aβ-EVs in axons (**Figures 5A^iii^** and **A^iv^**). We then questioned if each individual EV regulated intra-axonal protein synthesis. Thus, we measured puromycin incorporation in axons around the EV docking/internalization site (5 µm radius) and compared the signal to that of axons that did not interact with EVs within each chamber. We observed no differences in local protein synthesis between DMSO-EV-PKH26-stained axons and PKH26 unlabeled axons (**Figure 5B**). Importantly however, when axonal compartments were treated with Aβ-EVs, a significant increase in puromycin incorporation in axonal segments surrounding the EV internalization/docking site was detected (**Figure 5C**). These results confirm that Aβ-EVs positively regulate local translation in axons.

**Figure 5.**
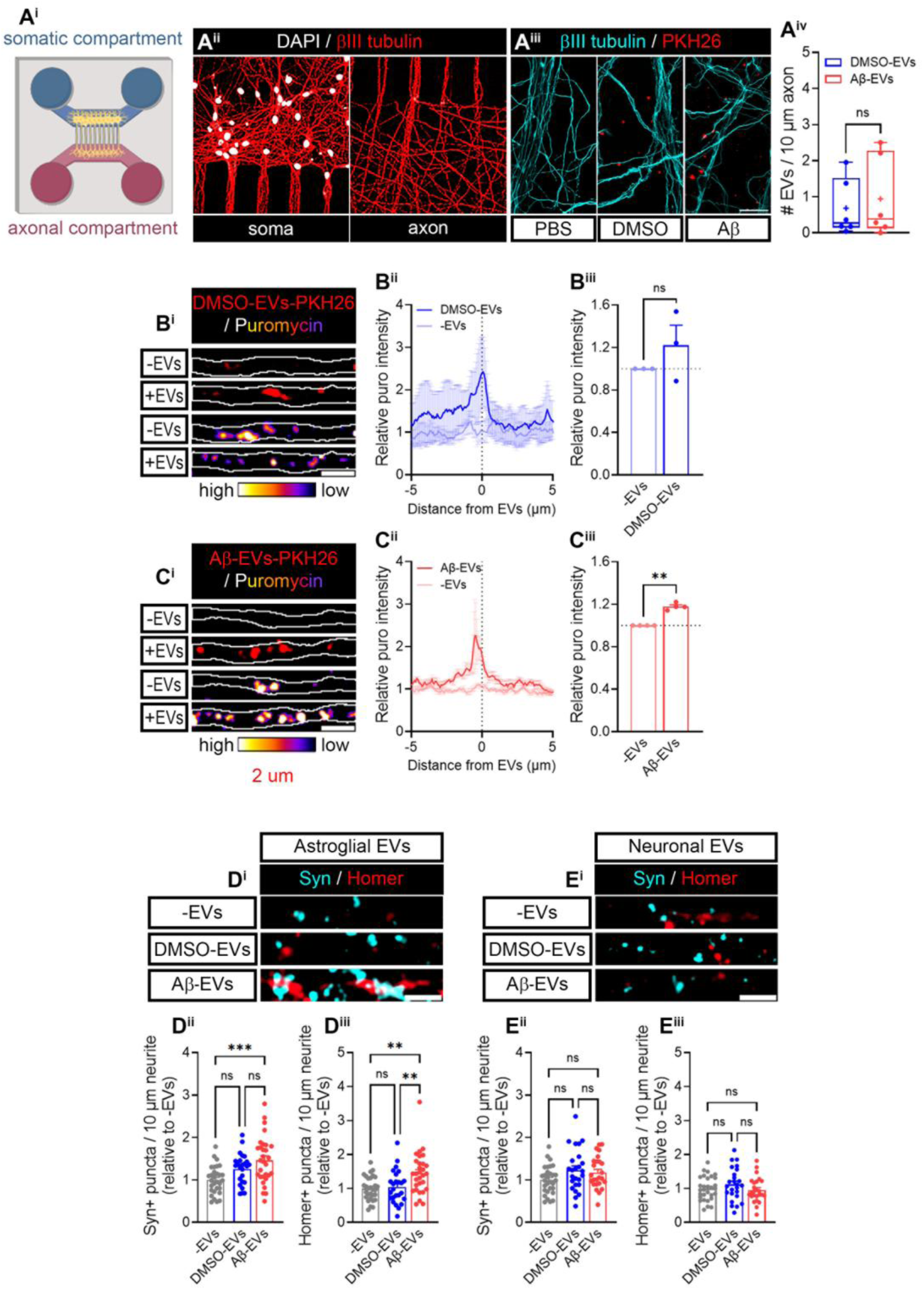
Aβ-EVs enhance local translation in axons and increase synaptic markers. **(A)** Microfluidic isolation of axons in cultured neurons. Schematic representation of a microfluidic chamber created in https://BioRender.com (**A^i^**). Neurons stained with an anti-βIII tubulin antibody and the nuclear marker DAPI show the fluidic isolation of the somatic compartment (left) and the axons (right) (**A^ii^**). Micrographs represent axons treated with PBS-PKH26 (negative control), DMSO-EVs-PKH26 or Aβ-EVs-PKH26. Scale bar = 25 µm (**A^iii^**). The box and whisker graph represents the number of DMSO-EVs or Aβ-EVs docked/internalized in axons (10 µm-long segments). Data was retrieved from 6 individual microfluidic chambers (n=6) seeded in 4 independent cultures (N=4). t-test; n.s, not.significant. **(B)** DMSO-EVs do not induce changes in local translation. Micrographs show non-EV- and DMSO-EV-associated axons (upper panels) and the corresponding puromycin labelling (represented as heatmaps; lower panels). Scale bar = 2 µm (**B^i^**). Linescans representing the average puromycin intensity ±SEM measured around the EV docking/internalization site (± 5 µm) relative to non-EV-associated axons measured in 3 individual chambers form 3 independent experiments (n=3; N=3) (**B^ii^**). The bar graph summarizes the data represented in the linescans. t-test; n.s, not significant (**B^iii^**). **(C)** Aβ-EVs enhance local protein synthesis in axons. Micrographs show non-EV- and Aβ-EV-associated axons (upper panels) and the corresponding puromycin labelling (represented as heatmaps; lower panels). Scale bar = 2 µm (**C^i^**). Linescans representing the average puromycin intensity ±SEM measured in 4 individual chambers around the EV docking/internalization site (± 5 µm) relative to non-EV-associated axons (**C^ii^**). The bar graph summarizes the data represented in the linescans. t-test; **p < 0.01 (**C^iii^**). **(D)** Astroglia Aβ-EVs lead to increased levels of synaptic markers in neurons. Micrographs show the levels of presynaptic marker synaptophysin (Syn) and post-synaptic protein Homer-1 (Homer) in neurites upon treatment with non-EV supplemented medium (upper panel), or media containing DMSO-EVs (middle panel) or Aβ-EVs (lower panel). Scale bar = 2 µm (**D^i^**). The bar graphs represent the mean number ±SEM of Syn+ (**D^ii^**) or Homer+ (**D^iii^**) puncta in neurites treated with DMSO- or Aβ-EVs relative to non-EV-treated neurons. Data corresponds to 25-30 neurites from 5 independent cultures (n=25-30; N=5). One-way ANOVA followed by Holm-Šídák *post hoc* test for selected pairs of columns. **p < 0.01; ***p < 0.001; n.s, not significant. **(E)** Neuronal EVs do not regulate synaptic markers. Micrographs show the levels of Syn and Homer in neurites upon treatment with non-EV supplemented medium (upper panel), or media containing DMSO-EVs (middle panel) or Aβ-EVs (lower panel). Scale bar = 2 µm (**E^i^**). The bar graphs represent the mean number ±SEM of Syn+ (**E^ii^**) or Homer+ (**E^iii^**) puncta in neurites treated with DMSO- or Aβ-EVs relative to non-EV-treated neurons. Data corresponds to 25-30 neurites from 5 independent cultures (n=25-30; N=5). One-way ANOVA followed by Holm-Šídák *post hoc* test for selected pairs of columns. n.s, not significant.

Whether the regulation of local translation by astroglia-derived EVs has a broader impact on neuronal function remains unclear. Notably, under physiological conditions, local translation in neurons is involved in axon pathfinding, guidance, arborization and maintenance, as well as in synapse formation and synaptic plasticity ^12–15^. However, this mechanism might become dysregulated in neurodegenerative diseases and lead to neuronal dysfunction ^37^. For instance, treatment of neurons with non-EV-associated Aβ peptides leads to widespread changes in the axonal transcriptome ^38^ and enhances local protein synthesis *in vitro* and *in vivo* ^16,35^. In this context, locally synthesized proteins have been involved in contributing to Alzheimer’s disease (AD) pathology ^16,17,35,39^. AD is characterized by synaptic dysfunction in early stages of the disease, and downregulation of synaptic markers has been reported in preclinical stages ^40^. Thus, we wondered if Aβ-EVs regulated synaptic proteins. We treated neurons with astroglial DMSO-EVs and Aβ-EVs for 24 hours and analyzed the presence of pre- and post-synaptic markers along neurites (synaptophysin and Homer-1 respectively). Unexpectedly, puncta containing synaptophysin or Homer-1 were significantly increased when neurons were treated with Aβ-EVs but not with DMSO-EVs, when compared to non-EV supplemented media (**Figure 5D**). Again, we used neuronal EVs as comparative controls and observed that neither DMSO-EVs nor Aβ-EVs enhanced synaptic markers (**Figure 5E**). We did however detect a trend towards an increase of synaptophysin in neurons treated with neuronal DMSO-EVs (which became significant when comparing non-EV supplemented media to DMSO-EV treatment through t-test; p=0.041), in line with previously reported data that indicate that neuronal EVs released in basal conditions enhance synaptic protein levels ^41^. All in all, these data suggest that astroglial Aβ-EVs not only upregulate local neuronal translation, but they also enhance levels of synaptic proteins. Based on our findings we then asked if both effects of astrocyte-derived Aβ-EVs were mechanistically linked. Thus, we next performed proteomic analyses to identify astroglial EV cargoes that could explain both phenotypes.

### 5. Aβ-induced vesicular Rps6 secretion by astrocytes enhances local protein synthesis and increases synaptic markers in neurons

We subjected astrocyte-derived EVs to mass spectrometry and analyzed their protein cargo. Background peptide levels were established by performing proteomic analyses on the same serum-free medium in which cells were cultured. We observed the enrichment of 391 proteins present in astroglial EVs regardless of the experimental condition (**Figure 6A^i^**). Once again, neuronal EVs were used for comparison, which showed 431 enriched proteins compared to the culture medium (**Figure 6A^ii^**). Venn diagrams indicated that, whereas 287 proteins were detected in both astroglial and neuronal EVs, 104 were exclusively released by astrocytes and 144 by neurons (**Figure 6A^iii^**). We then performed GO term analyses to identify BPs and signaling pathways significantly represented in EVs. BP clustering indicated that proteins found in both astroglia and neuronal EVs were mainly involved in translation regulation. However, from the 10 most enriched clusters, 6 of them contained astroglia vesicular proteins related to this molecular mechanism, while translation regulators found in neuronal EVs were represented in only 4 enriched clusters (**Figure 6B^i^**). These results suggest that astroglia EVs regulate protein synthesis in acceptor cells more likely than neuronal EVs. Indeed, the largest cluster found in glial EVs based on KEGG pathways analysis was that containing ribosomal proteins (**Figure 6B^ii^**). Our previous results had demonstrated that ribosomal proteins were overall upregulated in neuritic extracts isolated from neuron-astrocyte co-cultures compared to neuronal monocultures (**Figure 2B**). Thus, we focused on the presence of ribosomal proteins in EVs. It is noteworthy to comment that eukaryotic ribosomes contain ca. 80 ribosomal proteins, and that 31 of them were found in neuronal EVs while 45 were present in astroglial EVs. Thus, more than 50% of proteins contained in ribosomes were detected in vesicles released by astrocytes. Venn diagrams indicated that 28 ribosomal proteins were commonly found in both neuronal and glial EVs, while 3 of them were only detected in vesicles released by neurons, and 17 were exclusively found in astroglial EVs (**Figure 6C^i^**). Interestingly, amongst these 17 proteins we identified Rps6, whose levels we had seen enhanced in neuron-astrocytes co-cultures upon exposure to Aβ oligomers (**Figure 2D**). Thus, we focused on vesicular Rps6. By further inspecting the proteomic data we observed that this ribosomal protein was significantly enriched in astroglial Aβ-EVs, compared to the culture medium, but not in DMSO-EVs (**Figure 6C^ii^**). The presence of Rps6 in vesicular extracts from Aβ-treated astrocytes was later confirm by western blotting (**Figure 6D**).

**Figure 6.**
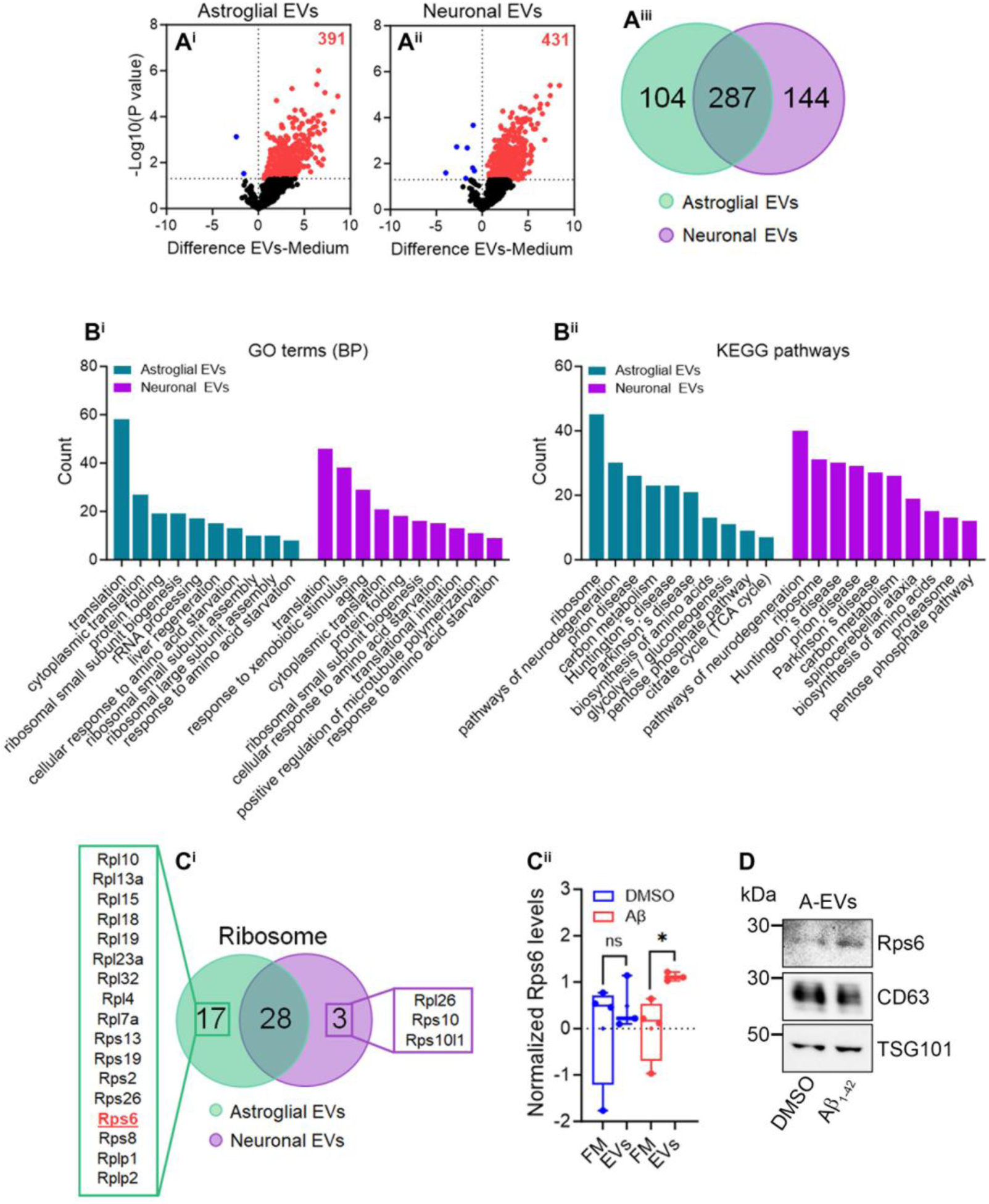
Astroglia EVs are enriched in ribosomal proteins. **(A)** Protein enrichment in astroglia and neuronal EVs identified by mass spectrometry. The volcano plots represent the upregulation (red dots) and downregulation (blue dots) of proteins detected in vesicular extracts from astrocytes (**A^i^**) or neurons (**A^ii^**) compared to serum-free culture medium, regardless of the treatment with vehicle or Aβ oligomers. Results are the average of 3 independent samples from astrocytes (N=3) and 4 samples from neurons and serum-free medium (N=4). Venn diagram depicting the overlap of proteins found enriched in astroglia and neuronal EVs (**A^iii^**). **(B)** Translation regulators are enriched in astroglia EVs. The bar graphs represent the functional clustering, identified with DAVID, of glial (green bars) and neuronal (purple bars) EV cargoes based on the biological process (**B^i^**) or the pathways they are involved in (**B^i^**^i^). Only the 10 most represented clusters are plotted (p < 0.05). **(C)** Ribosomal protein Rps6 is present at detectable levels in astroglia Aβ-EVs. Venn diagram depicting the overlap of ribosomal proteins found enriched in astroglia and neuronal EVs. The identity of 17 and 3 proteins exclusively found in astroglia and neuronal EVs respectively is shown (**C^i^**). Box and whisker graphs indicate the normalized levels of ribosomal protein Rps6 found in serum-free media treated with vehicle or Aβ (4 samples; N=4) and in vesicular extracts from astrocytes (3 samples; N=3) by proteomics. Pairwise t-tests comparing EVs to serum-free medium in control or Aβ-induced conditions. *p < 0.05; n.s, not significant (**C^ii^**). Western blot showing the presence of Rps6 in astroglia vesicular extracts (**C^iii^**).

Next, we sought to determine if astrocyte-released Rps6 was directly involved in local translation in neurons. To that end, we genetically downregulated astroglial *Rps6* with siRNAs (**Supplementary Figure 4B**). siRNA-transfected astrocytes were then treated with vehicle or Aβ oligomers and EVs were isolated from the culture medium by ultracentrifugation. EVs, labelled with PKH26, were then applied to axons grown in microfluidic chambers. Similar to previous experiments, we qualitatively observed a larger amount of PKH26-labelled Aβ-EVs that control EVs regardless of the transfection of astrocytes with control- of *Rps6* siRNA (**Figure 7A^i^**). However, after a 45-minute treatment no changes were detected in the docking/internalization within axons between experimental conditions (**Figure 7A^ii^**). We then analyzed puromcycin incorporation in axons (as a measurement for local protein synthesis) around the EV docking/internalization site and compared the signal to that of axons that did not interact with EVs within each chamber. Consistent with our previous observations, DMSO-EVs did not enhance local translation in axons, neither when astrocytes were treated with control siRNA nor with *Rps6* siRNA (**Figures 7B^i^**, **7B^ii^**and **7B^v^)**. Conversely, puromycin incorporation increased in axon segments close to the docking/internalization sites of Aβ-EVs released by control-transfected astrocytes, while this effect was blocked if axons were treated with Aβ-EVs isolated from glia transfected with *Rps6* siRNA (**Figures 7B^iii^-7B^iv^**). These results confirm that local neuronal translation is regulated by vesicular Rps6 released by astrocytes exposed to Aβ oligomers.

**Figure 7.**
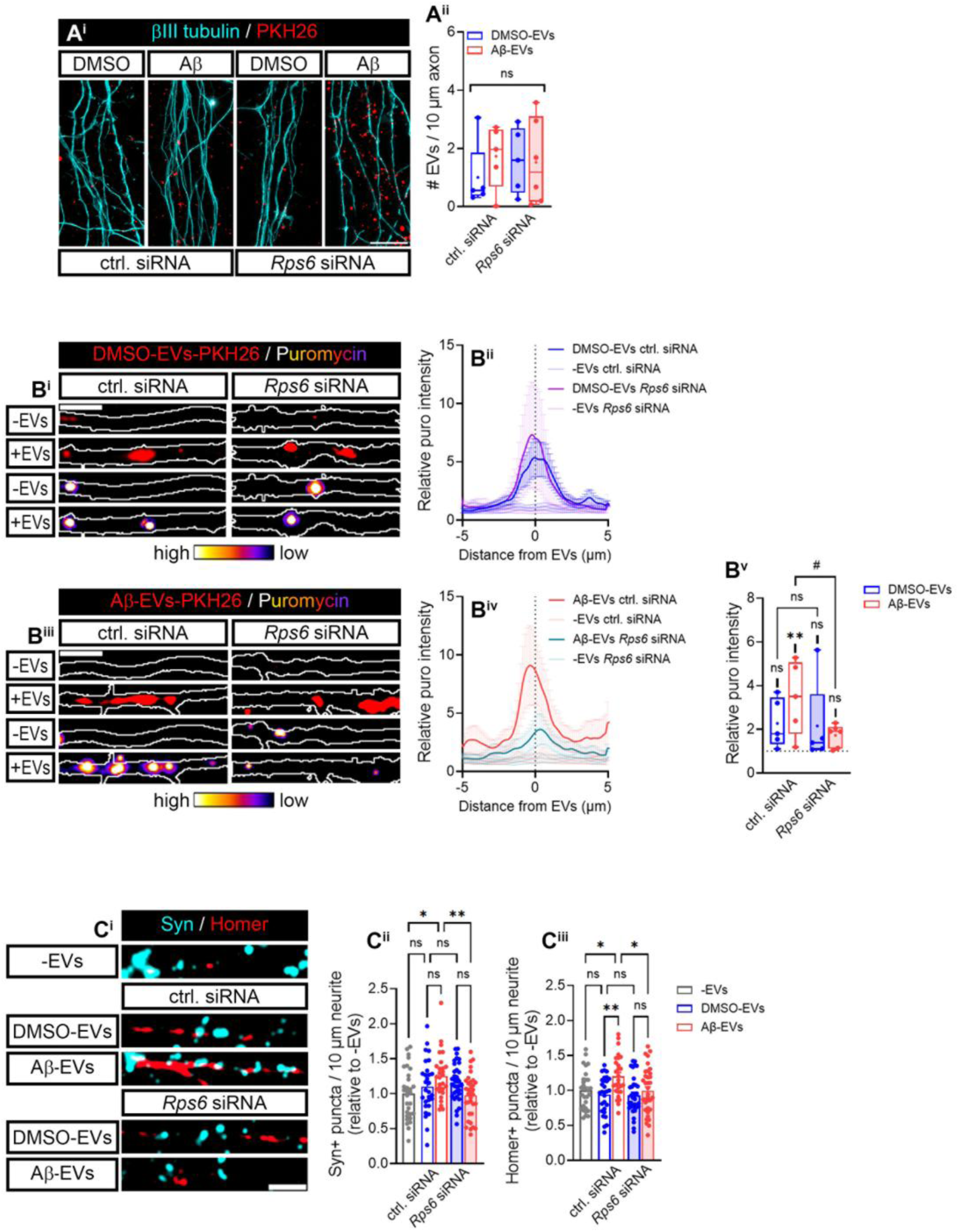
Vesicular Rps6 released by Aβ-treated astrocytes regulates local translation and synaptic markers in neurons. **(A)** *Rps6* knockdown in astrocytes does not affect EV docking to and/or internalization within axons. Micrographs show axons treated with DMSO-EVs-PKH26 or Aβ-EVs-PKH26 isolated from astrocytes transfected with control- or *Rps6* siRNAs. Scale bar = 25 µm (**A^i^**). The box and whisker graph represents the number of DMSO-EVs or Aβ-EVs docked/internalized in axons (10 µm-long segments) in each experimental condition described in (**A^i^**). Data was retrieved from 5-6 individual microfluidic chambers (n=5-6) seeded in 3 independent cultures (N=3). One-way ANOVA; n.s, not.significant (**A^ii^**). **(B)** Astroglia Aβ-EV-dependent local translation in axons is blocked by *Rps6* knockdown. Micrographs representing non-EV- and DMSO-(**B^i^**) or Aβ-EV-associated axons (**B^iii^**) and the corresponding puromycin intensity (heatmaps) are shown. DMSO- and Aβ-EVs were isolated from control- and *Rps6* siRNA-treated astrocytes as indicated. Scale bar = 2 µm. Linescans represent the average puromycin intensity ±SEM measured around the EV docking/internalization site (± 5 µm) relative to non-EV-associated axons. EVs were isolated from control (**B^ii^**) and Aβ-treated astrocytes (**B^iv^**) transfected with a control or an *Rps6* siRNA as indicated. Analyses were performed in 5-6 individual chambers form 3 independent cultures (n=5-6; N=3). The box and whisker graph summarizes the data represented in the linescans shown in (**B^ii^**) and (**B^iv^**). One-way ANOVA followed by Holm-Šídák *post hoc* test was performed. The asterisks indicate significant differences detected between non-EV associated axons and those axons associated with EVs released in each experimental condition: **p < 0.01; n.s, not significant. The hash sign represents significant changes in local protein synthesis detected in axons treated with different EVs. #p < 0.05; n.s, not significant (**B^v^**). **(C)** Aβ-EV-dependent regulation of synaptic markers is mediated by astroglia-derived Rps6. Micrographs show the levels of Syn and Homer in neurites upon treatment with non-EV supplemented medium, or media containing DMSO-EVs or Aβ-EVs released by control- or *Rps6* siRNA-transfected astrocytes. Scale bar = 2 µm (**C^i^**). The bar graphs represent the mean number ±SEM of Syn+ (**C^ii^**) or Homer+ (**C^iii^**) puncta in neurites treated with DMSO- or Aβ-EVs relative to non-EV-treated neurons under the experimental conditions described in (**C^i^**). Data corresponds to 28-35 neurites from 4 independent cultures (n=28-35; N=4). One-way ANOVA followed by Holm-Šídák *post hoc* test for selected pairs of columns. *p < 0.05; **p < 0.01; n.s, not significant.

Finally, we wanted to address if astroglial Rps6-mediated local protein synthesis affected the levels of synaptic markers (**Figure 7C**). Thus, we again transfected astrocytes with control- or *Rps6* siRNAs and exposed them to vehicle or Aβ for 24 hours. Astrocyte-derived vesicles when then used to treat neurons for 24 hours and levels of pre- and post-synaptic markers (synaptophysin and Homer-1, respectively) were measured by immunocytochemistry. No changes were observed when neurons were treated with DMSO-EVs regardless of glia transfection with control- or *Rps6* siRNA. On the other hand, like in previous results Aβ-EVs released by control-transfected astrocytes enhanced the number of puncta containing synaptophysin (**Figures 7C^i^** and **7C^ii^**) or Homer (**Figure 7C^i^** and **7C^iii^**) compared to non-EV supplemented medium. Importantly, positive puncta were significantly reduced when Aβ-EVs were released by *Rps6*-transfected astrocytes. Thus, neuronal local translation modulation by astroglial Rps6 leads to increased levels of synaptic markers.

Overall, our results suggest a model in which astrocyte-derived EVs released in response to Aβ contain ribosomal proteins, such as Rps6, that regulate local translation in neurons through non-cell-autonomous mechanisms, whereby locally synthesized proteins enhance synaptic markers potentially contributing to synaptic function (**Figure 8**).

**Figure 8.**
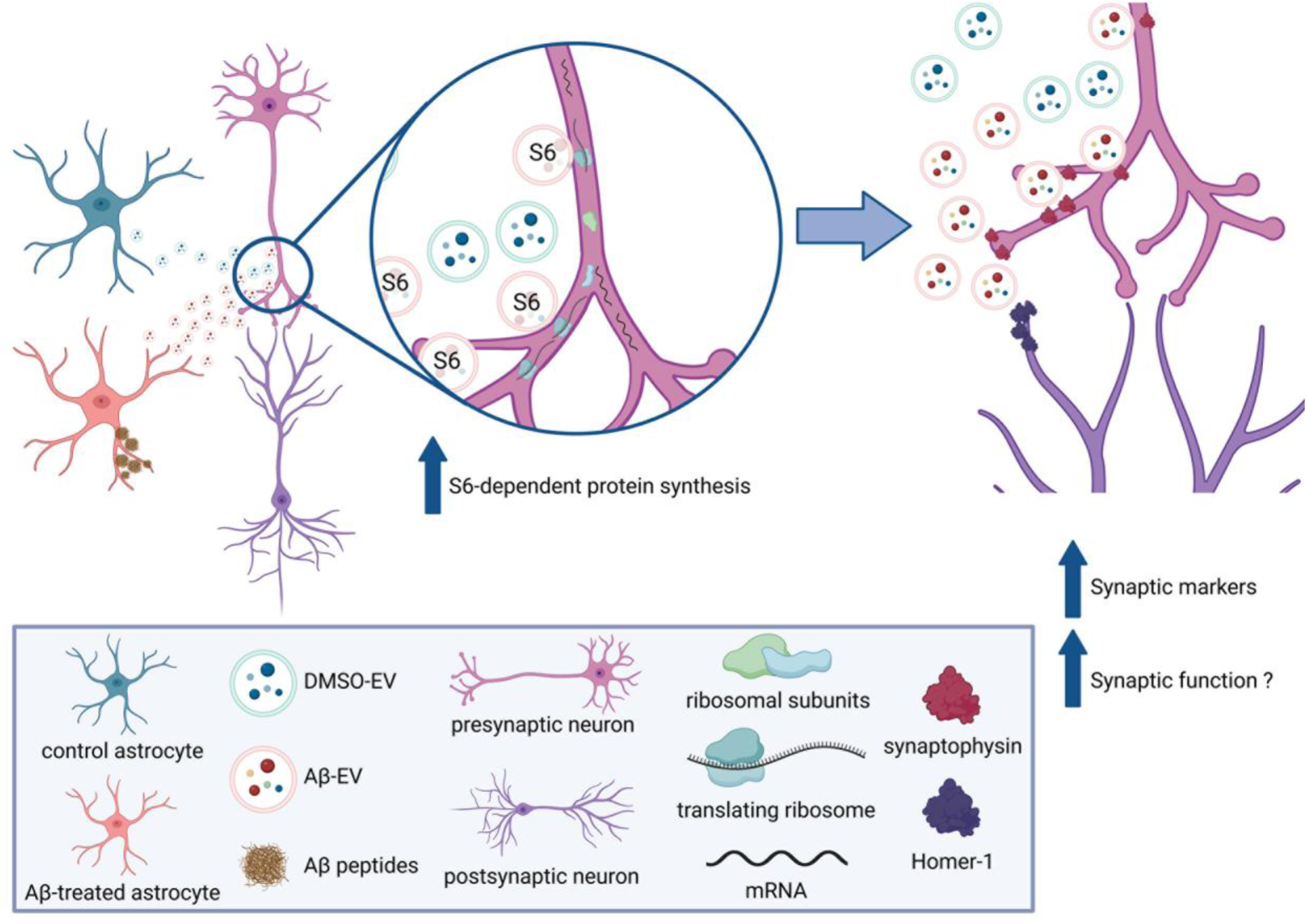
Proposed model for astroglia Aβ-EV-induced local translation and synaptic marker regulation in neurons. Non-chronical exposure of astrocytes to Aβ peptides induces the release of EVs containing translation regulators (including components of the ribosomes). Ribosomal proteins (specifically Rps6) are transferred from glia to neurons thus modulating local translation in the latter. Locally synthesized proteins in neurons, whose levels depend on Rps6 released by astroglia Aβ-EVs, in turn enhance synaptic markers, potentially contributing to synaptic function.

## Discussion

Local translation enables neurites to rapidly respond to extracellular cues, a process essential for proper nervous system function. Indeed, disruption of mRNA localization and/or translation within subneuronal compartments has been increasingly involved in the progression of neurodegenerative diseases, including AD ^37^. Within this context, *in vitro* exposure of axons to Aβ peptides, key mediators of AD pathology, has shown to induce protein synthesis in axons, contributing to neurodegeneration ^16,17^. However, whether local translation is entirely regulated by the neuron itself or if it can be modulated by neighboring glial cells, such as astrocytes, remains largely unexplored.

Although recent evidence has shown that astrocyte-secreted factors induce *de novo* protein synthesis in synaptic compartments ^42^, no previous data indicate that astrocytes modulate neuronal local translation through EVs. While Schwann cells in the PNS can transfer ribosomes and RNAs to injured axons to promote regeneration ^16,19–21^, the existence of a similar mechanism in the CNS has not been established.

Since Aβ treatment is known to increase axonally synthetized proteins ^16,35^, we first assessed whether Aβ exposure enhances protein synthesis within neurites from neuron monocultures or neuron-astrocyte co-cultures. On the one side, OPP-biotin conjugation in neuritic extracts (Figure 1B) revealed increased levels of newly synthesized proteins in neurites after Aβ treatment. However, somewhat unexpectedly, results from desomatized neurons could not conclusively distinguish whether these proteins were locally synthesized or transported from the soma. This discrepancy with prior reports may result from differences in culture systems (microfluidic devices vs. Boyden chambers), protein labeling methods (fluorescent non-canonical amino acids vs. OPP tagging), or even the source of primary neurons (hippocampal vs. cortico-hippocampal). OPP-biotin labeled polypeptides were detectable in neurites in neuron-astrocyte co-cultures in both basal and amyloid induced conditions (**Figure 1B^ii^**). More importantly, we found that OPP-labeled proteins were still detectable in neurites devoid from somatic input only when co-cultured with astrocytes and treated with Aβ (**Figure 1B^iii^**), indicating that under these conditions, astrocytes support local protein synthesis in neurons in response to amyloid treatment.

To investigate this further, proteomic analysis of neuritic extracts from neuronal monocultures and neuron-astrocyte co-cultures revealed that astrocytes modulate a distinct subset of proteins enriched in translational machinery, including ribosomal proteins, and translation initiation and elongation factors (**Figure 2A and B^i^**). Among these, astrocytes significantly upregulated ribosomal proteins Rpl26 and Rps6, components of the 60S and 40S ribosomal subunits, respectively, in neurites following Aβ exposure (**Figure 2B^ii^**). These findings were validated by immunocytochemistry (**Figure 2C and D**). Rpl26 has previously been detected in axons of different neuronal types under basal conditions ^20,43^. Moreover, *Rpl26* local translation has been detected within postsynaptic compartments of *in vitro* synaptosomes upon direct exposure to astrocyte-conditioned medium ^42^. As for Rps6, this ribosomal protein localizes near the mRNA and tRNA binding sites within the ribosome and is therefore considered a key player in translation activation ^44^. Rps6 has been linked to regeneration in both the PNS and the CNS. For instance, Rps6 phosphorylation increases following sciatic nerve injury and is associated with axonal regenerative capacity ^45,46^. Similarly, Duan and colleagues reported elevated Rps6 levels in retinal neurons after axotomy, which were associated with mTOR signaling and neuronal survival ^47^. Rps6 is also localized to axons upon Aβ exposure, coinciding with increased intra-axonal protein synthesis, which in this case leads to neuronal death rather than survival ^16^. Interestingly, the Rps6/mTOR signaling pathway is one of the best-characterized mechanisms involved in local translation regulation ^48,49^. Therefore, it is reasonable to speculate that astrocytes may increase axonal Rps6 levels in response to Aβ to influence local protein synthesis and potentially drive neurons toward either survival or death likely through mTOR signaling pathway.

We next explored the regulatory mechanism by which astrocytes influence local protein synthesis in neurites. Given the non-contact setup in Boyden chambers, we hypothesized that cell-to-cell communication is mediated by secreted factors. Among potential mediators, EVs emerged as strong candidates. Particles secreted by astrocytes and neurons were characterized according to the ISEV guidelines, ^50^ and we confirmed their identity as EVs through multiple experimental approaches. In particular, cryo-EM revealed that our nanoparticle preparations are enriched in membrane-bound vesicles, consistent with their classification as EVs. Astrocyte- and neuron-derived EVs were characterized according to ISEV guidelines ^50^ and confirmed their identity via multiple experimental approaches. Morphological differences were observed between neuron- and astrocyte-derived EVs (**Figure 3B**), further supporting distinct functional roles. Although no specific EV subpopulation (microvesicles, exosomes or apoptotic bodies) was distinguished, from cryo-EM and NTA results we can conclude that at least astroglial EVs likely consist of a mixed population of microvesicles and exosomes. Moreover, we found that Aβ exposure significantly increased EV secretion, particularly from astrocytes (**Figure 3C-D**).

Our results demonstrate for the first time the contribution of astrocytic EVs to neuronal local translation, since astrocyte-derived EVs released in response to Aβ act as positive regulators of intra-axonal protein synthesis (**Figure 4A** and **Supplementary figure 3B**). These results were further confirmed in neurons treated with astroglia conditioned medium in which EV secretion was impaired by *Rab11A* downregulation (**Figures 4B** and **4C**) and in neurons exposed to EV-depleted medium (**Supplementary figure 3C**).Importantly, neuron-derived EVs did not replicate these results (**Figure 4B** and **Supplementary figure 3D**, confirming the specificity of astroglia vesicles in neuronal local translation regulation.

To definitively assess local effects of astroglia Aβ-EVs exclusively in axons, we used microfluidic chambers to isolate axons from somata in a spatial and fluidically isolated manner. In these devices, astroglial-derived EVs were observed docked at/or internalized within axons (**Supplementary Figure 4A** and **Figure 5A^iii^**). In all experiments the amount on EVs used for treatments was normalized to the number of secreting cells. Given that Aβ enhances EV release by astrocytes, it is possible that a greater abundance of Aβ-EVs exerts a stronger effect compared to DMSO-EVs on neurons. Thus, in microfluidic chamber we addressed the effect of individual PKH-26-labelled EV on axons and confirmed that local translation increased at axonal sites in contact with Aβ-EVs. This effect was not observed when the same strategy was conducted with DMSO-EVs. In sum, these findings reinforce the conclusion that EVs released by astrocytes in response to Aβ drive intra-axonal translation in neurons.

Beyond local translation, we explored the impact of Aβ-EVs on synaptic compartments. Previous studies have linked astrocyte-derived EVs to both neuroprotective and neurotoxic effects, depending on the context. For instance, astroglial EVs have been reported to contain pro-survival factors such as synapsin or neuroglobin ^4,51^ or to enhance neurite arborization in anti-inflammatory induced conditions ^52^. Conversely, other authors found that astrocyte-derived EVs induced inflammation and neuronal death in amyloid pathology models ^53–55^. In our study, astrocyte-derived Aβ-EVs significantly increased puncta containing the pre- and post-synaptic markers synaptophysin and Homer-1, respectively, along neurites, suggesting a protective effect (**Figure 5D**). On the other hand, neuron-derived DMSO-EVs marginally increased synaptophysin in neurites, consistent with previous reports suggesting that neuronal EVs sustain synaptic protein levels ^41^ (**Figure 5E**).

Following the findings that on the one hand Aβ-EVs modulate local protein synthesis and on the other hand they increase synaptic protein levels, we next demonstrated a mechanistic link between both effects. First, we found that astroglial EVs contain more than 50 % of all existing ribosomal proteins in mammalian cells (**Figure 6C^i^**). More importantly, Rps6 was specifically enriched in astrocyte-derived Aβ-EVs (**Figures 6C** and **6D**). This finding is consistent with the upregulation of Rps6 in axons when neurons were cultured in the presence of astrocyte (**Figure 2**) Knockdown of *Rps6* in astrocytes abolished the axonal response to astroglial Aβ-EVs in terms of both translation regulation and synaptic markers enhancement (**Figures 7C and 7D**). These results suggest that EV-mediated delivery of Rps6 in response to amyloid is required for astroglial modulation of local protein synthesis and synaptic integrity. Traditionally, synaptic function and plasticity have been associated with postsynaptic translation ^12,56–58^. However, growing evidence underscores the relevance of intra-axonal translation in synapse formation and synaptic activity ^59–62^. . Our findings support this notion, proposing that astrocyte-derived EVs serve as vehicles for delivering ribosomal components, including Rps6, to axons where they enhance local translation which in turn leads to synaptic stability (**Figure 8**).

To date, astroglial-derived EVs have been involved in functions such as neurite outgrowth and arborization, or apoptosis regulation, without further exploring the mechanisms underlying these responses. Here we describe, for the first time, a novel mechanism of intercellular communication in which astrocytes respond to Aβ exposure by releasing EVs enriched in ribosomal proteins. These EVs are taken up by axons, where they stimulate local protein synthesis and promote synaptic marker expression, unravelling a previously unknown neuroprotective role for astroglial EVs. These findings contribute to understanding how glial cells influence neuronal function and open new avenues for therapeutic strategies targeting glia-neuron communication in AD and related disorders.

## Author contributions

<u>María Gamarra</u>: experimental design, methodology, formal analysis, writing (original draft), manuscript review and editing. <u>Aida de la Cruz-Gambra</u>: methodology, formal analysis (synaptic marker quantification), writing (original draft), manuscript review and editing. <u>Maite Blanco-Urrejola</u>: methodology, formal analysis, manuscript review. <u>Esperanza González</u>: experimental design (EV isolation and characterization), methodology, formal analysis (nanoparticle tracking), manuscript review and editing. <u>Mikel Azkargorta</u>: experimental design (proteomics), formal analysis (proteomics), manuscript review. <u>Felix Elortza</u>: experimental design (proteomics), manuscript review and editing. <u>Juan Manuel Falcón-Pérez</u>: experimental design (EV isolation and characterization), manuscript review and editing. <u>Jimena Baleriola:</u> conceptualization, experimental design, methodology, formal analysis, funding acquisition, project management, writing (original draft), manuscript review and editing.

## Acknowledgements

We acknowledge the Electron Microscopy and Crystallography Platform from the Center for Cooperative Research in Biosciences (CIC bioGUNE) for assistance in the characterization of extracellular vesicles by cryo-EM.

## Data availability

The datasets generated for this study are available on request to the corresponding author.

## Supplementary figures

**Supplementary figure 1.**
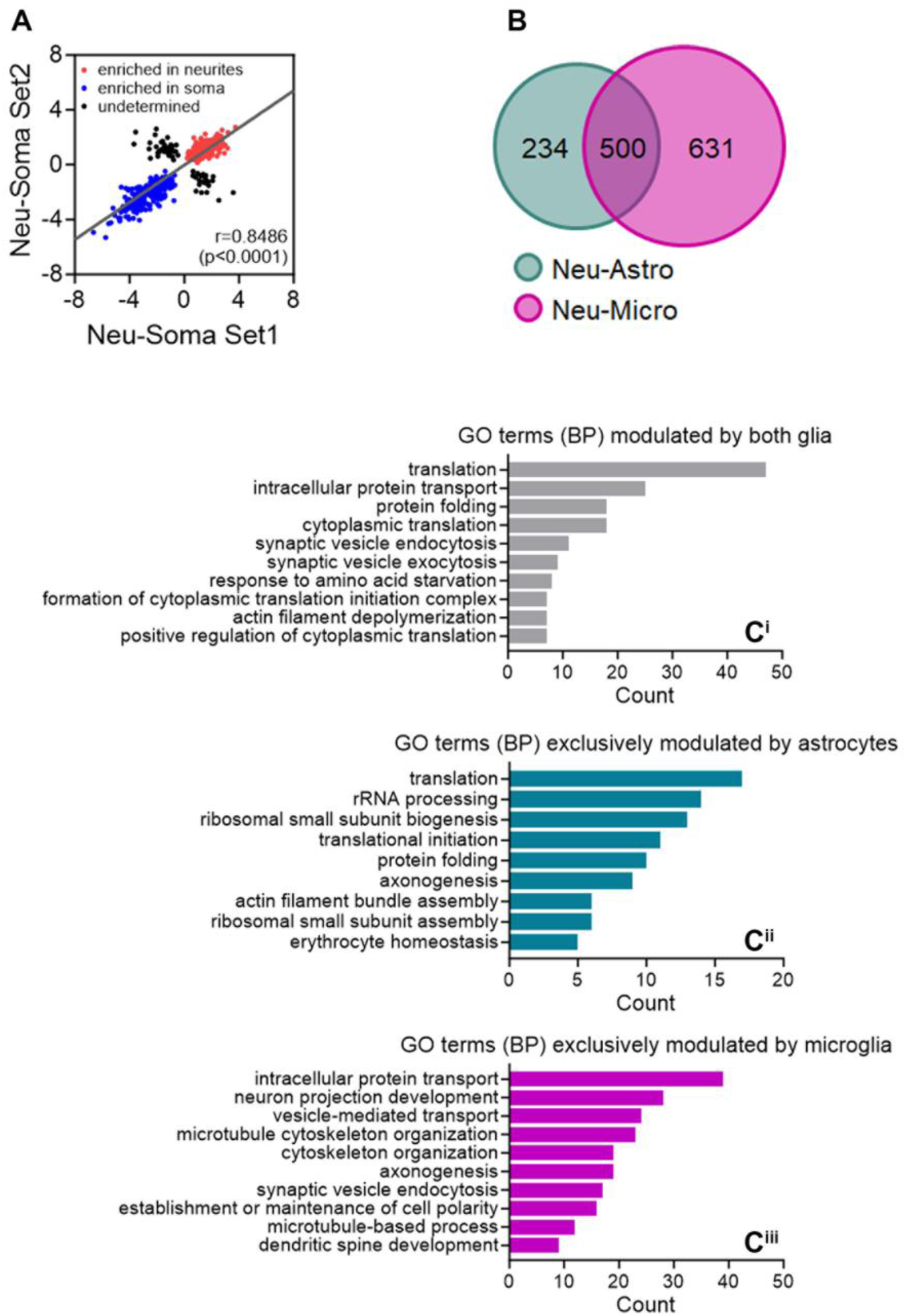
Neuritic proteins are selectively regulated by astrocytes or microglia. **(A)** Analyses of neuritic proteins commonly increased (red dots) or decreased (blue dots) in two unrelated datasets where proteomic analyses were performed in neuronal monocultures or glia-neuron co-cultures (namely neuron-astrocyte and neuron-microglia co-cultures). The graph indicates the correlation between proteins upregulated or downregulated in neurites compared to the soma in neuronal monocultures in both sets of experiments. A significant correlation was found between both datasets (p < 0.0001). Pearson r is also indicated. **(B)** Venn diagrams depicting the overlap of proteins regulated in astrocyte-neuron co-cultures or microglia-neuron co-cultures compared to neuronal monocultures. Proteins whose fold-change in neurites compared to the soma in neuronal cultures were discordant among datasets were discarded (black dots in **(A)**). **(C)** Astrocytes and microglia regulate common but also distinct protein in neurites. Functional clustering performed with DAVID (based on the biological process) identified proteins whose levels were changed by both the presence of astroglia and microglia in neuronal cultures (**C^i^**), as well as those regulated either by astrocytes (**C^ii^**) or microglia (**C^iii^**).

**Supplementary figure 2.**
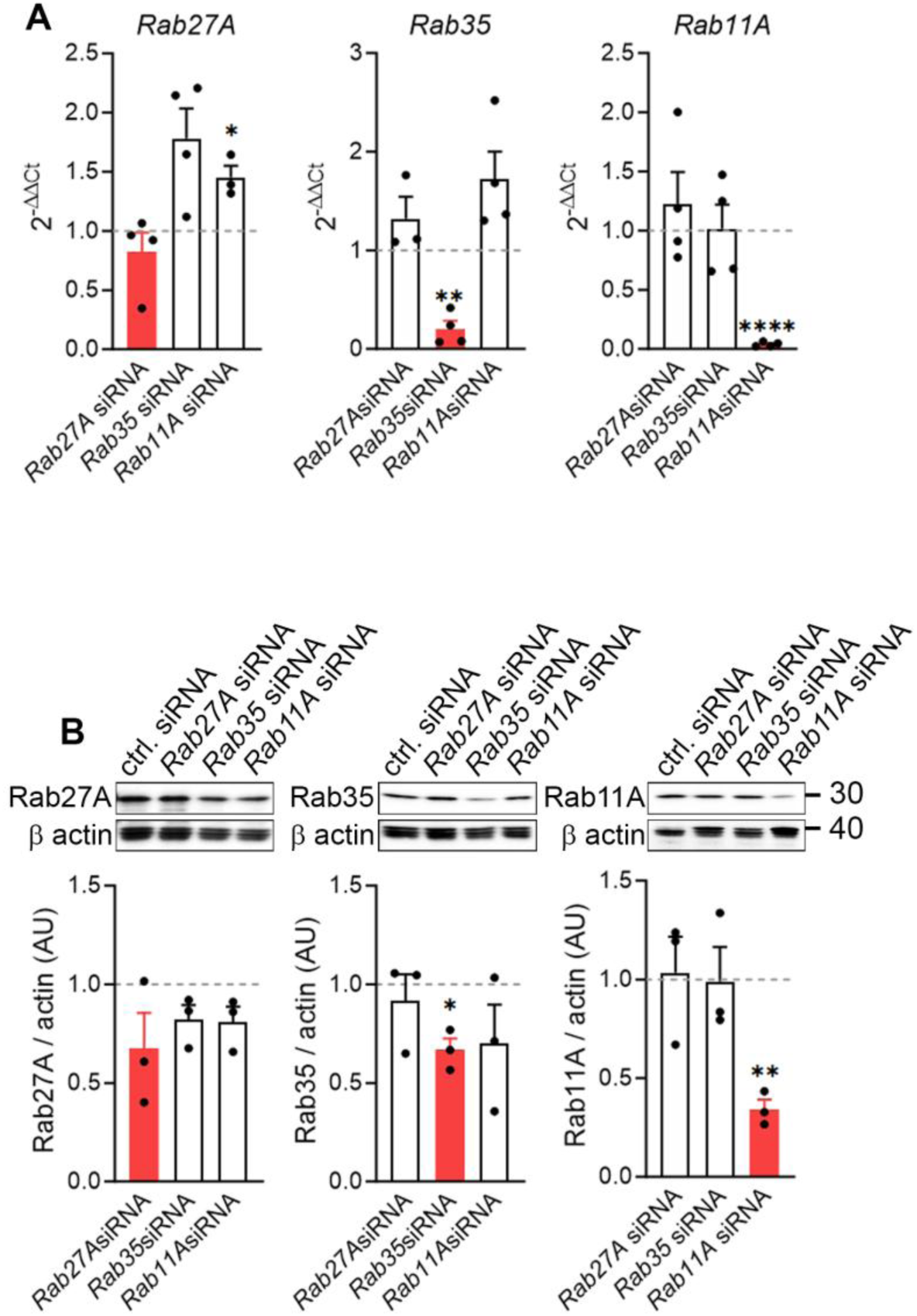
Knockdown efficiency of *Rab27A*, *Rab35* and *Rab11A* in astrocytes. **(A)** Quantitative RT-PCR analyses of *Rab27A*, *Rab35* and *Rab11A* mRNAs upon transfection of astrocytes with the corresponding siRNAs compared to control-transfected cells. One-way ANOVA analysis with Dunnet’s *post hoc* tests was performed to compare the effect of specific siRNAs to control-transfected cells. *p < 0.05; **p < 0.01; ****p < 0.0001. **(B)** Western blot analyses of Rab27A, Rab35 and Rab11A protein upon transfection of astrocytes with the corresponding siRNAs compared to control-transfected cells. One-way ANOVA analysis with Dunnet’s *post hoc* tests was performed to compare the effect of specific siRNAs to control-transfected cells. *p < 0.05; **p < 0.01.

**Supplementary figure 3.**
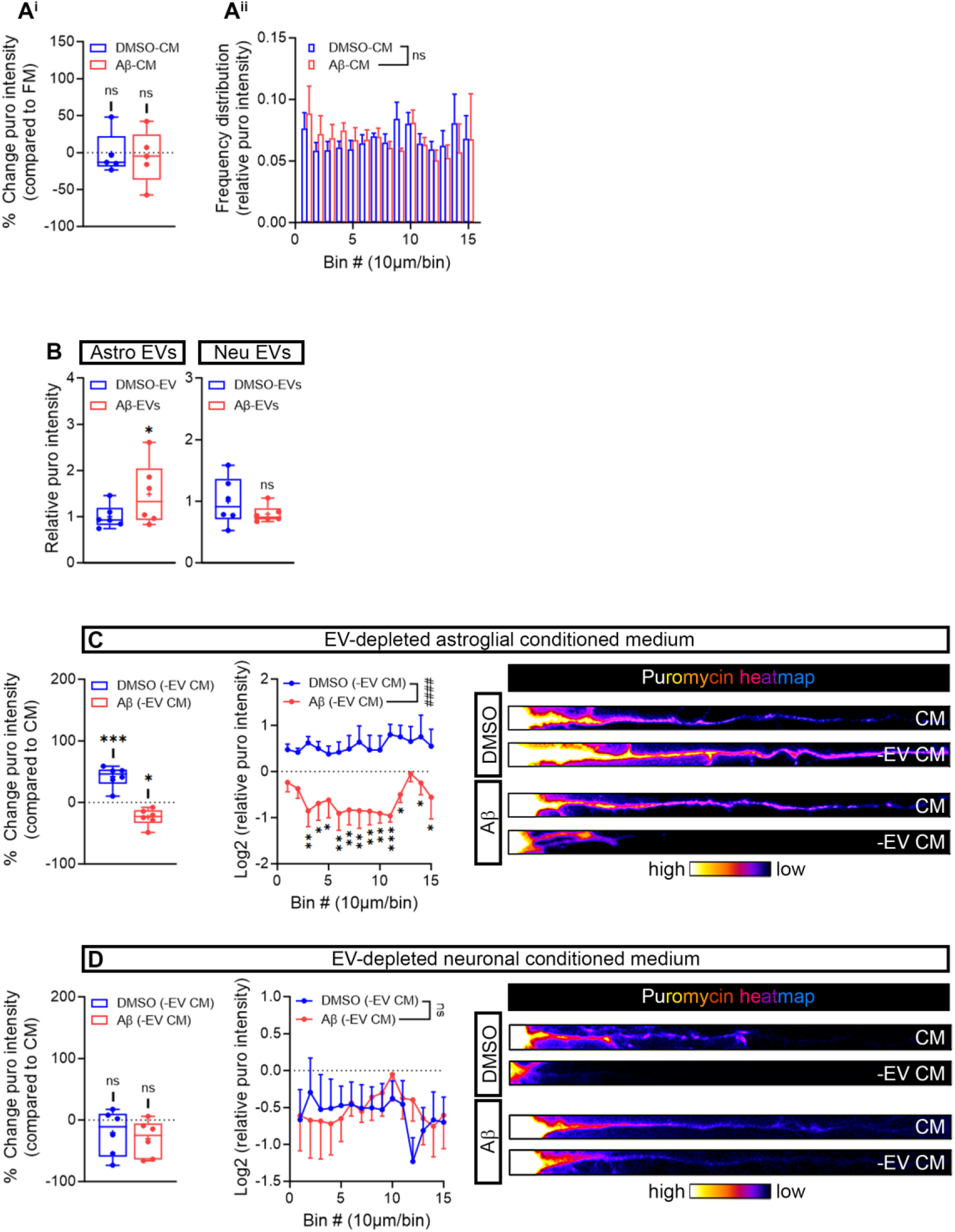
Effect of astroglia conditioned medium or EV-depleted astroglia conditioned medium on newly-synthesized axonal proteins. **(A)** Conditioned medium released by control or Aβ-treated astrocytes does not affect puromycin levels in axons after a 45-minute treatment. Box and whisker graph representing percent changes in puromycin fluorescent intensity in axons from conditioned media (CM) released by DMSO- or Aβ-treated astrocytes compared to fresh medium-treated neurons (FM). One-way ANOVA followed by Dunnet’s *post hoc* test comparing the effect of DMSO- or Aβ-CM to FM. n.s, not significant (**A^i^**). Intensity vs distance graph representing puromycin frequency distribution in axons measured in 15 10-µm bins covering a distance of 150 μm from the soma. Neurons were treated with conditioned media from control astrocytes or Aβ-treated astrocytes. Data corresponds to 5 independent experiments (N=5). Two-way ANOVA: n.s, not significant (**A^ii^**). **(B)** Astroglia Aβ-EVs enhance protein synthesis after a 24 h-treatment. Box and whisker graphs representing the relative puromycin fluorescent intensity in astroglial (left) or neuronal (right) DMSO-EV- and Aβ-EV-treated neurons measured in 6 independent cultures (N = 6). t-test: *p < 0.05: n.s; not significant. **(C)** EV depletion from culture media confirms the positive effect of astroglia Aβ-EVs on newly synthesized axonal medium. The box and whisker graph represents percent changes in puromycin fluorescent intensity in neurons treated with media depleted from DMSO-EVs or Aβ-EVs with respect to their corresponding complete media (CM). One-way ANOVA followed by Dunnet’s *post hoc* test comparing the effect of each EV-depleted media to CM. ***p < 0.001; *p < 0.05 (**C**; left graph). Intensity vs distance graph representing puromycin intensity in axons measured in 15 10-µm bins covering a distance of 150 μm from the soma. Data are plotted as puromycin intensity relative to the CM (Log2 was applied for better visualization in the graph). Two-way ANOVA followed by Holm-Šídák *post hoc* test. The hash signs indicate overall changes detected by treatment with both EV-depleted media. ####p < 0.0001. Asterisks depict changes between media depleted from DMSO-EVs and Aβ-EVs at specific distances from the soma. *p < 0.05; **p < 0.01; ***p < 0.001. Statistical analyses were performed in results retrieved from 6 independent experiments (N=6) (**C**, right graph). **(D)** Neuronal EV depletion from culture media does not affect puromycin labelling in axons. The box and whisker graph represents percent changes in puromycin fluorescent intensity in neurons treated with media depleted from DMSO-EVs or Aβ-EVs with respect to their corresponding complete media (CM). One-way ANOVA followed by Dunnet’s *post hoc* test comparing the effect of each EV-depleted media to CM. n.s, not significant (**D**; left graph). Intensity vs distance graph representing puromycin intensity in axons measured in 15 10-µm bins covering a distance of 150 μm from the soma. Data are plotted as puromycin intensity relative to the CM (Log2 was applied for better visualization in the graph). Two-way ANOVA followed by Holm-Šídák *post hoc* test. n.s, not significant. Statistical analyses were performed in results retrieved from 6 independent experiments (N=6) (**D**; right graph).

**Supplementary figure 4.**
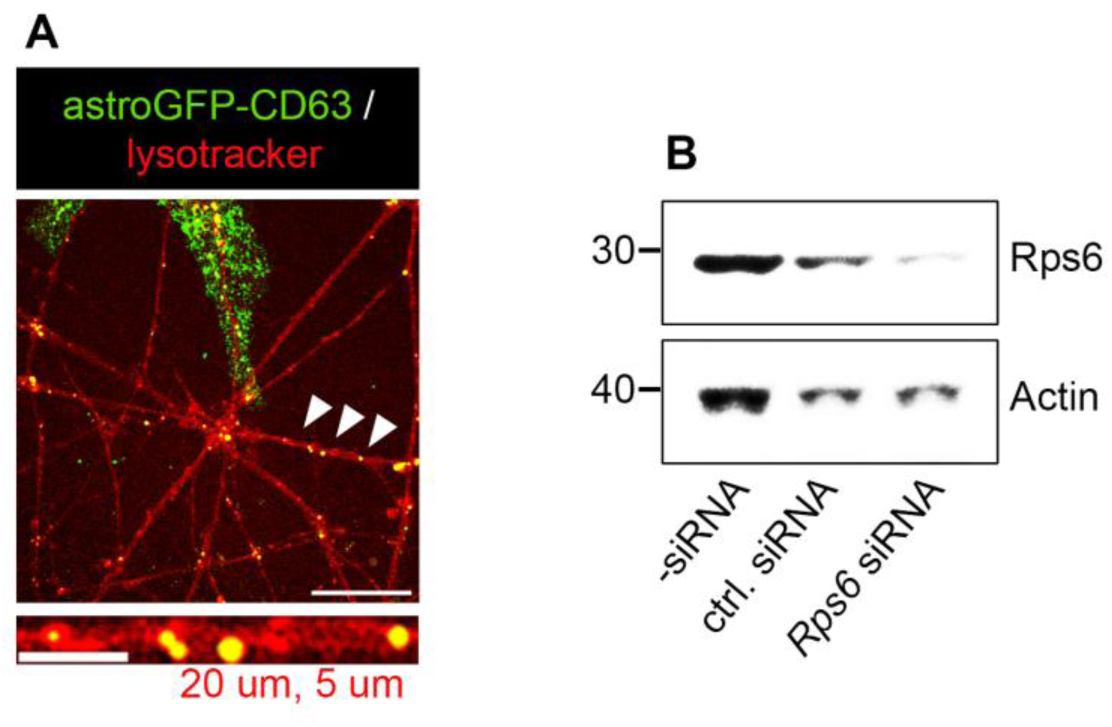
**(A)** Confirmation of EV transfer from astrocytes to axons in microfluidic chambers. Astrocytes were transfected with myrGFP-CD63 controlled by GFAP promoter. Transfected astrocytes were cultured in the axonal compartment of microfluidic chambers myrGFP-tagged vesicle were visualized in axons. Scale bar = 20 µm (5 µm in inset). **(B)** Downregulation of Rps6 in astroglia lysates following transfection of a targeting siRNA.

